# Efficacy of a Broadly Neutralizing SARS-CoV-2 Ferritin Nanoparticle Vaccine in Nonhuman Primates

**DOI:** 10.1101/2021.03.24.436523

**Authors:** Michael G. Joyce, Hannah A. D. King, Ines Elakhal Naouar, Aslaa Ahmed, Kristina K. Peachman, Camila Macedo Cincotta, Caroline Subra, Rita E. Chen, Paul V. Thomas, Wei-Hung Chen, Rajeshwer S. Sankhala, Agnes Hajduczki, Elizabeth J. Martinez, Caroline E. Peterson, William C. Chang, Misook Choe, Clayton Smith, Parker J. Lee, Jarrett A. Headley, Mekdi G. Taddese, Hanne A. Elyard, Anthony Cook, Alexander Anderson, Kathryn McGuckin-Wuertz, Ming Dong, Isabella Swafford, James B. Case, Jeffrey R. Currier, Kerri G. Lal, Robert J. O’Connell, Sebastian Molnar, Manoj S. Nair, Vincent Dussupt, Sharon P. Daye, Xiankun Zeng, Erica K. Barkei, Hilary M. Staples, Kendra Alfson, Ricardo Carrion, Shelly J. Krebs, Dominic Paquin-Proulx, Nicos Karasavva, Victoria R. Polonis, Linda L. Jagodzinski, Mihret F. Amare, Sandhya Vasan, Paul T. Scott, Yaoxing Huang, David D. Ho, Natalia de Val, Michael S. Diamond, Mark G. Lewis, Mangala Rao, Gary R. Matyas, Gregory D. Gromowski, Sheila A. Peel, Nelson L. Michael, Diane L. Bolton, Kayvon Modjarrad

## Abstract

The emergence of novel severe acute respiratory syndrome coronavirus 2 (SARS-CoV-2) variants stresses the continued need for next-generation vaccines that confer broad protection against coronavirus disease 2019 (COVID-19). We developed and evaluated an adjuvanted SARS-CoV-2 Spike Ferritin Nanoparticle (SpFN) vaccine in nonhuman primates (NHPs). High-dose (50 *µ*g) SpFN vaccine, given twice within a 28 day interval, induced a Th1-biased CD4 T cell helper response and a peak neutralizing antibody geometric mean titer of 52,773 against wild-type virus, with activity against SARS-CoV-1 and minimal decrement against variants of concern. Vaccinated animals mounted an anamnestic response upon high-dose SARS-CoV-2 respiratory challenge that translated into rapid elimination of replicating virus in their upper and lower airways and lung parenchyma. SpFN’s potent and broad immunogenicity profile and resulting efficacy in NHPs supports its utility as a candidate platform for SARS-like betacoronaviruses.

**One-Sentence Summary:** A SARS-CoV-2 Spike protein ferritin nanoparticle vaccine, co-formulated with a liposomal adjuvant, elicits broad neutralizing antibody responses that exceed those observed for other major vaccines and rapidly protects against respiratory infection and disease in the upper and lower airways and lung tissue of nonhuman primates.

---

The coronavirus disease 2019 (Covid-19) pandemic, caused by severe acute respiratory syndrome coronavirus 2 (SARS-CoV-2), has reached a milestone with the emergency use authorization and increasing availability of efficacious vaccines (*1*). Successes in rapid coronavirus vaccine development, however, have been tempered by the rise of virus variants (*2*). The accelerating frequency with which variants are emerging raises the prospect that host selective pressures may be driving evolution of mutants to escape vaccine-elicited immunity (*3*). This concern, coupled with stringent cold-chain requirements for product stability and high unit costs (*4, 5*), justifies the continued development of cost-effective, thermo-stable vaccines that match current ones in safety and efficacy, but provide broader coverage against a wide range of circulating variants and evolving strains, as well as novel species that may arise from zoonotic reservoirs in the future.

Self-assembling protein nanoparticle vaccines offer the advantage of multivalent antigen presentation, a property previously shown to augment immunogenicity over monovalent immunogens (*6-8*). Ferritin is a naturally occurring, ubiquitous, iron-carrying protein that self-oligomerizes into a 24-unit spherical particle (*9*). The three-fold axis symmetry of the resulting polymer makes it conducive to conjugation and antigen display of trimeric glycoproteins, such as SARS-CoV-2 Spike (S). Ferritin has been evaluated as a vaccine platform for several pathogens (*10-12*)—most notably influenza, for which it has demonstrated immune potency and breadth (*13, 14*). As such, ferritin vaccines have advanced to phase 1 clinical trials as a strategy to target multiple influenza strains (*15, 16*).

The prefusion-stabilized form of S is the basis for most major SARS-CoV-2 vaccine candidates (*17, 18*). Although a correlate of protection from Covid-19 has not been conclusively defined, there is mounting evidence that neutralizing, and some fraction of non-neutralizing, antibodies against S are necessary, if not sufficient, to confer protective immunity (*19, 20*). The most potent neutralizing antibodies are directed against the S receptor-binding domain (RBD), which mediates attachment to the primary host cell receptor, ACE-2. Prior assessment of a SARS-CoV-2 S ferritin nanoparticle (SpFN) vaccine candidate—co-formulated with a liposomal adjuvant (*21*)—has demonstrated potent immunogenicity and SARS-CoV-2 protection in mouse models (*unpublished*). These data provide a basis for evaluating SpFN immunogenicity and efficacy against viral replication and pathology in the airways and lungs of nonhuman primates (NHP), a standard model for preclinical evaluation of SARS-CoV-2 vaccines (*22*).

## Nanoparticle Vaccine and Study Design

The Spike Ferritin Nanoparticle (SpFN) vaccine was designed as a ferritin-fusion recombinant protein for expression as a nanoparticle. Briefly, the Spike (S) protein sequence was derived from the Wuhan-Hu-1 genome sequence (GenBank accession number: MN908947.3). The S ectodomain was modified to introduce two proline residues (K986P, V987P) and removal of the furin cleavage site (RRAS to GSAS), as previously described (*17*). To stabilize S trimer formation on the ferritin molecule, the heptad repeat between hinge 1 and 2 (residues 1140 – 1161) was mutated to stabilize coiled-coil interactions. An adjuvant—Army Liposomal Formulation QS21 (ALFQ)—was mixed with the SpFN vaccine at room temperature within four hours before administration. ALFQ formulation has been described previously (*23*). Briefly, it comprises dimyristoyl phosphatidylcholine (DMPC), dimyristoyl phosphatidylglycerol (DMPG), cholesterol (Chol), and synthetic monophosphoryl lipid A (3D-PHAD^®^) (Avanti Polar Lipids, Alabaster, AL) and QS-21 (Desert King, San Diego, CA).

In this study, 32 male and female specific-pathogen-free, research-naïve Chinese-origin rhesus macaques (age 3 - 7 years) were distributed—on the basis of age, weight and sex—into 4 cohorts of 8 animals (table *S1*). Animals were vaccinated intramuscularly with either 50 or 5 µg of SpFN, formulated with ALFQ, or 1ml of phosphate buffer solution (PBS) in the anterior proximal quadriceps muscle, on alternating sides with each dose in the series. Immunizations were administered twice—4 weeks apart—or once, 4-weeks prior to challenge (fig. *S1*). Animals were challenged with 1×10^6^ TCID50 of SARS-CoV-2 (BEI Resources, NIAID, NIH: SARS-Related Coronavirus 2, Isolate USA-WA1/2020, NR-53780 (Lot# 70038893) administered simultaneously by the intratracheal (1.0 ml) and intranasal (0.5 ml per nostril) route.

### Vaccine Immunogenicity

#### Serum antibody responses

We measured longitudinal antibody responses in animals after each vaccination and viral challenge by the Meso Scale Discovery (MSD) electrochemiluminescence platform. Total binding to SARS-CoV-2 prefusion stabilized S protein (S-2P) (*17*) increased from baseline to an area under the curve (AUC) of 679,213 and 1,646,288 at 4 weeks after two vaccinations with 5 and 50 µg of SpFN, respectively (Fig. 1A). Vaccination with a single 50 µg dose resulted in a 4-week AUC of 621,605. Binding responses were unchanged in vaccinated groups after viral challenge; whereas, unvaccinated controls had a 200-fold rise. Two-doses of 5ug or 50 µg SpFN elicited reciprocal 50% inhibitory dilution (ID50) neutralizing antibody geometric mean titers (GMT) of 22,405 and 52,773, respectively, 2 weeks after second vaccination, and leveled off at 12,171 and 22,527 2 weeks later (Fig. 1B). Single dose 50 µg SpFN elicited a peak GMT of 4063. Authentic virus neutralization activity mirrored group differences seen in the pseudovirus assay, but at somewhat lower values (Fig. 1C).

**Fig. 1.**
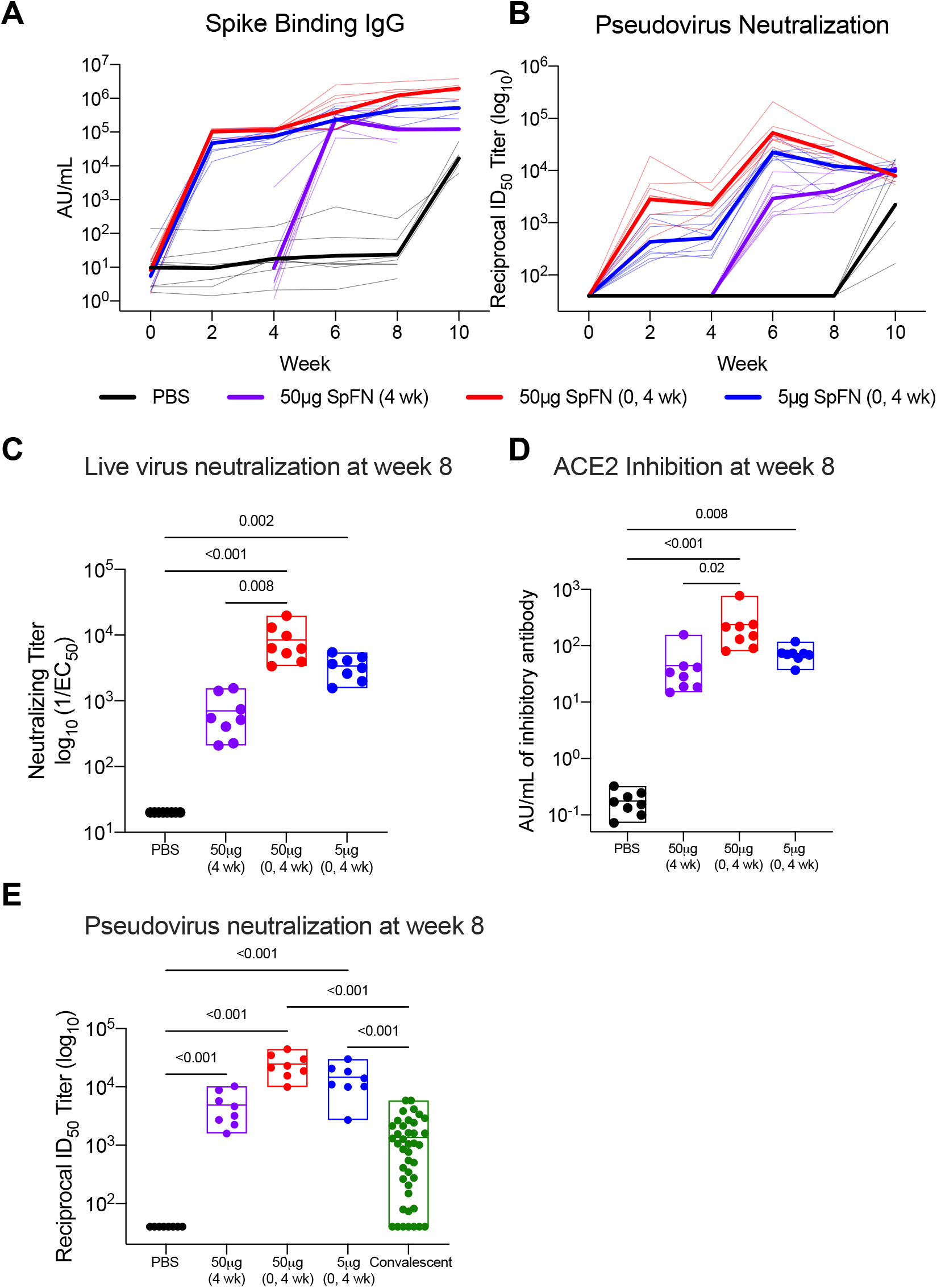
SpFN elicited binding and neutralizing antibody responses elicited in rhesus macaques. (**A, B**) Animals were vaccinated with 5 or 50 µg of SpFN at weeks 0 and 4 or 50 µg of SpFN at week 4 only. Control animals were given phosphate-buffered saline (PBS) instead. Serum specimens were assessed for (**A**) SARS-CoV-2 Spike-specific IgG by the MSD electrochemiluminescent platform and (**B**) SARS-CoV-2 pseudovirus neutralization every 2 weeks following vaccination and 1 to 2 weeks following viral challenge. Data are depicted as the area under the curve (AUC) of IgG binding and virus neutralization reciprocal 50% inhibitory dilution (ID_50_), respectively. Thick lines indicate geometric means within each group and thin lines represent individual animals. (**C**) Authentic virus neutralization was also assessed at 4 weeks after last vaccination. (**D**) Inhibition of angiotensin-converting enzyme 2 (ACE2) receptor binding to the receptor-binding domain (RBD) at 4 weeks after last vaccination was measured on the MSD platform and reported in arbitrary units (AU)/ml. (**E**) Pseudoneutralization activity was compared to a panel of human convalescent sera (N=41 samples). In the box plots, horizontal lines indicate the mean and the top and bottom reflect the minimum and maximum. Symbols represent individual animals and overlap with one another for equal values where constrained. Significance was assessed using a Kruskal-Wallis test followed by a Dunn’s post-test.

We performed functional assessments of antibody responses by measuring the ability of sera to inhibit binding of RBD to the ACE2 receptor. Binding inhibition in the 5 and 50 µg vaccinated animals exceeded unvaccinated controls by a factor of 224 and 998, respectively (Fig. 1D). ACE2 competition in the single 50 µg dose group was 291 times higher than controls. We compared humoral responses to the vaccine against a panel of convalescent plasma samples with same pseudovirus neutralization assay; we found that two doses of either SpFN dose elicited neutralizing activity that was an order of magnitude higher than that of the convalescent sera (p<0.01) (Fig. 1E).

We used orthogonal approaches to assess binding antibody specificities to the Spike S1 subunit domains. RBD and S-2P binding, by MSD, recapitulated results of the ACE2 binding inhibition assay (fig. *S2*). Serum binding to the N-terminal domain (NTD), which may be a marker of additional protection through both neutralizing activity and non-neutralizing functions (*24*), were 500-fold higher compared to baseline, across vaccine groups (fig. *S3*). We assessed the strength of RBD binding by biolayer interferometry, finding an increasing antibody on-rate association response throughout follow-up (fig. *S4*). Given the potential importance of auxiliary antibody functions for protection (*25, 26*), we assessed a suite of Fc-mediated antibody effector functions, including opsonization, ADCD, ADCP, ADNP and trogocytosis (a measure of antigen transfer) (*27*). All activity peaked at week 6 and was highest in the two-dose 50 µg SpFN group (fig. *S5*).

#### Cellular immune responses

The character of the helper CD4+ T cell (Th) response is important for respiratory virus vaccine development, given the theoretical concern and precedent for vaccine-associated enhanced respiratory disease and its association with a Th2-biased response (*28*). We focused our assessment of cell-mediated immunity on canonical cytokines expressed by Th1 (interferon-gamma (IFN-γ), tumor necrosis factor (TNF), interleukin 2 (IL-2)) and Th2 (IL-4, IL-13) CD4 T cells. Robust Th1 responses were observed 4 weeks after second vaccination in all vaccinated groups, except one animal in the 50 µg single-dose group (Fig. 2A). Th1 responses were polyfunctional and variable but consistently high at week 8, ranging from 0.2% to 17%. Th2 and CD8+ T cell responses were minimal or undetectable (Fig. 2B, fig. *S6A*), though Th1 and Th2 responses correlated strongly (r=0.72, p<0.0001).

**Fig. 2.**
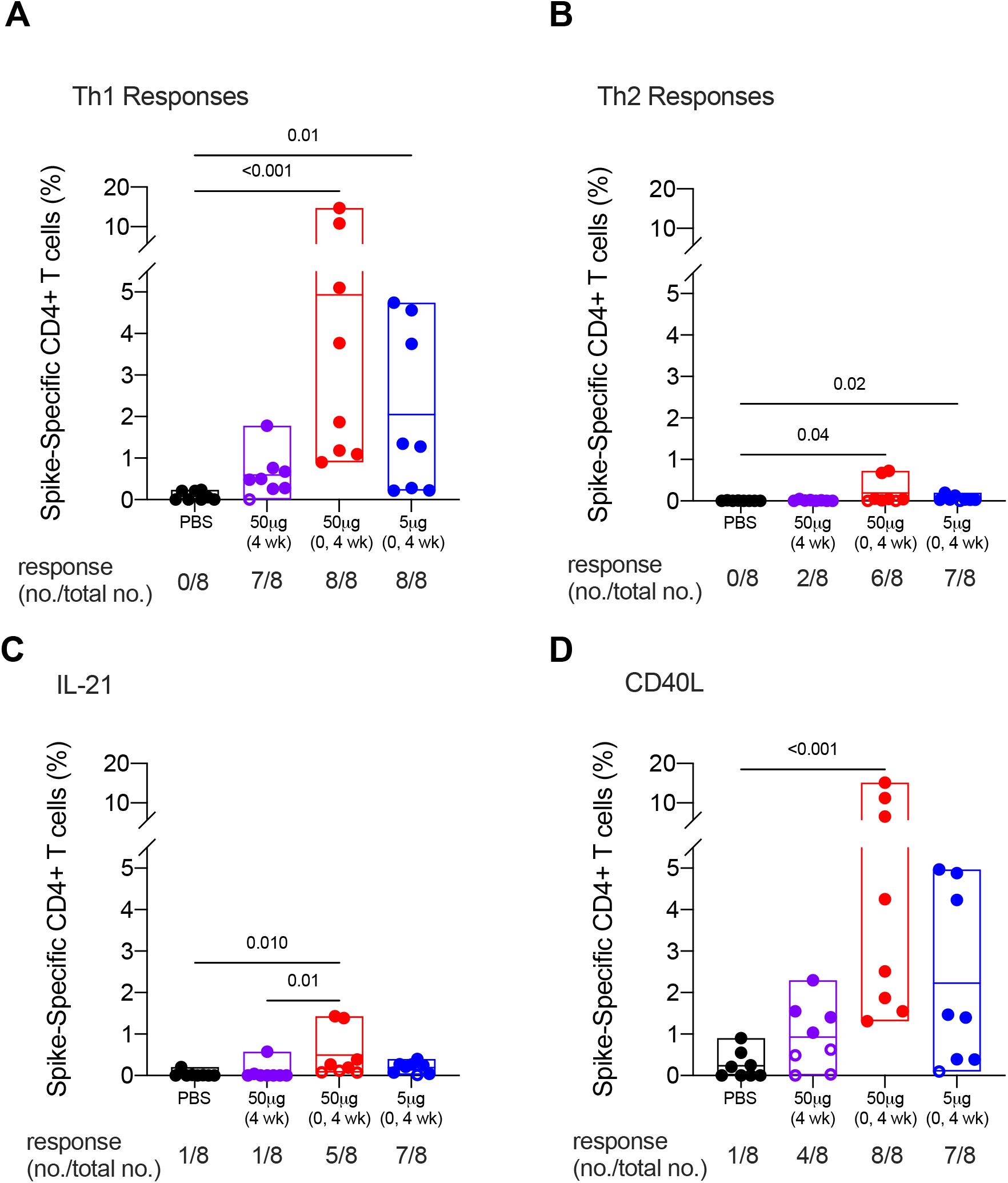
SpFN vaccine elicited SARS-CoV-2 Spike-specific CD4+ T cell responses in rhesus macaques. T cell responses were assessed by SARS-CoV-2 S peptide pool stimulation and intracellular cytokine staining of peripheral blood mononuclear cells collected at 4 weeks after last vaccination. S-specific memory CD4+ T cells expressing the indicated markers are shown as follows: (**A**) Th1 cytokines (IFNγ, TNF and IL-2); (**B**) Th2 cytokines (IL-4 and IL-13); (**C**) IL-21; and (**D**) CD40L. Boolean combinations of cytokine positive memory CD4+ T cells were summed. Probable positive responses, defined as >3 times the group background at baseline, are depicted as closed symbols. Positivity rates within each group are shown below each graph as a fraction. In the box plots, horizontal lines indicate the mean and the top and bottom reflect the minimum and maximum. Significance was assessed using a Kruskal-Wallis test followed by a Dunn’s post-test.

We interrogated key indicators of an engaged memory response such as IL-21, a cytokine, secreted by follicular helper CD4+ T cells (Tfh), that regulates the evolution of memory B cells (*29*). Five of eight animals dosed twice with 50 µg SpFN had IL-21 responses, as did seven of eight animals given 5 µg of vaccine (Fig. 2C). We also examined levels of CD40L: a broad T cell activation marker, expressed on the surfaces of CD4+ T cells and Tfh cells, that promotes B cell maturation through antibody isotype switching.(*29*) All but one animal receiving two-doses of SpFN had detectable CD40L+ responses (Fig. 2D), indicating an engaged memory response.

### Protection against high-dose SARS-CoV-2 respiratory challenge

#### Virologic Efficacy

Rhesus macaques generally exhibit mild disease that does not recapitulate the severe pneumonia observed in many people with COVID-19 (*22*). Protective efficacy, therefore, was assessed virologically and pathologically. The primary virologic endpoint was subgenomic mRNA copies/ml—an indicator of viral replication—in the upper (nasopharyngeal (NP) swabs, saliva) and lower airways (bronchoalveolar lavage (BAL) fluid) of vaccinated compared to control animals. On the second day after simultaneous respiratory challenge, sgmRNA levels in the BAL fluid of control animals peaked at a mean of 10^6^ copies/ml (Fig. 3A). In contrast, none of the eight animals that received two doses of 50 µg SpFN had detectable sgmRNA at day 2. By day 4 sgmRNA was undetectable in the BAL fluid of all animals of the 5 µg vaccine group and all but one animal that received single SpFN.

**Fig. 3.**
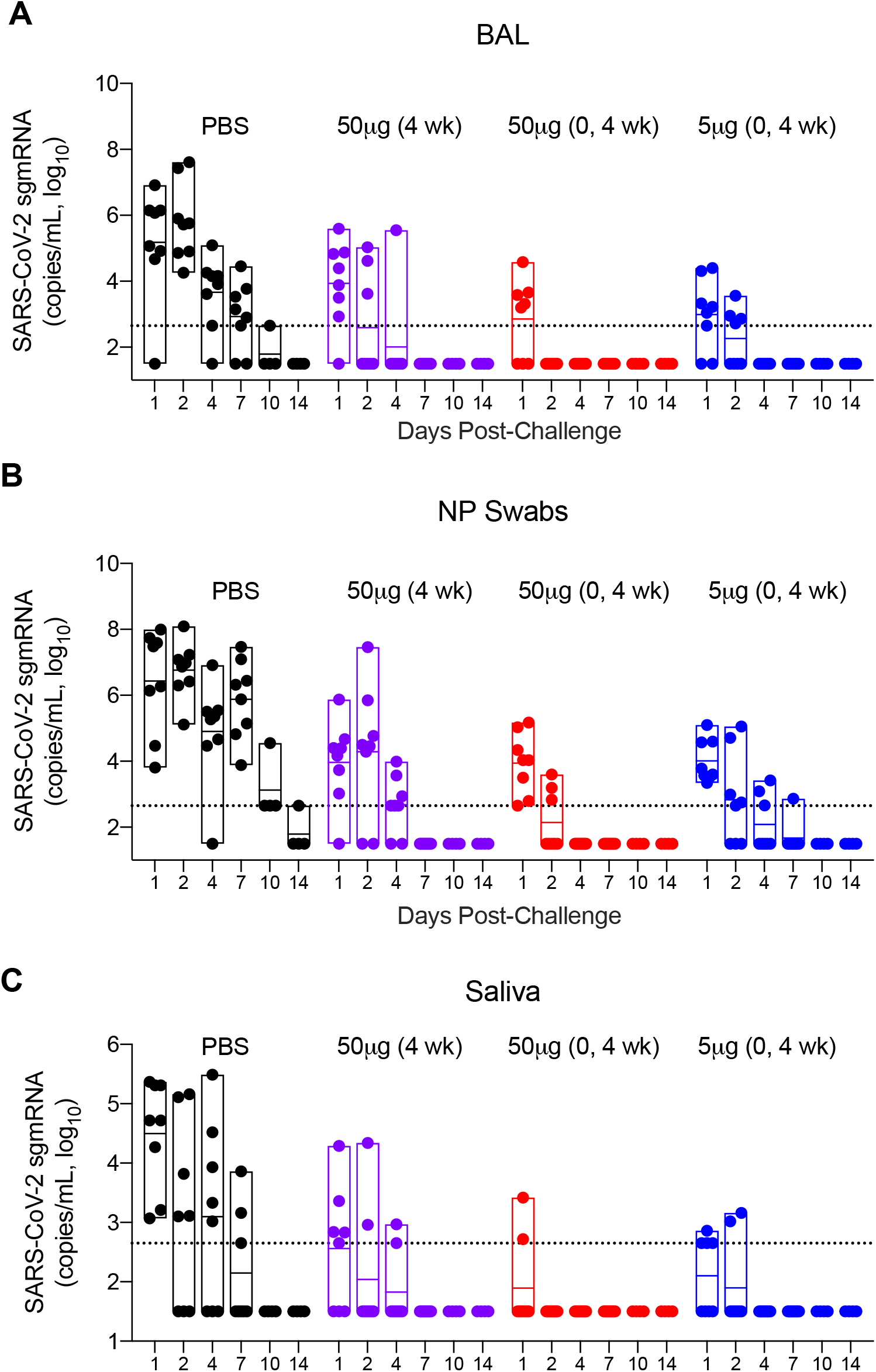
Viral replication in the lower and upper airways after SpFN vaccination and subsequent SARS-CoV-2 respiratory challenge. Subgenomic messenger RNA (sgmRNA) copies per milliliter were measured in the: (**A**) bronchoalveolar lavage fluid, (**B**) nasopharyngeal swabs and (**C**) saliva of vaccinated and control animals for two weeks following intranasal and intratracheal SARS-CoV-2 (USA-WA1/2020) challenge of vaccinated and control animals. Specimens were collected on 1, 2, 4, 7, 10 and 14 days post-challenge. Dotted lines demarcate assay lower limits of linear performance range (Log_10_ of 2.65 corresponding to 450 copies/ml). In the box plots, horizontal lines indicate the mean and the top and bottom reflect the minimum and maximum.

**Fig. 4.**
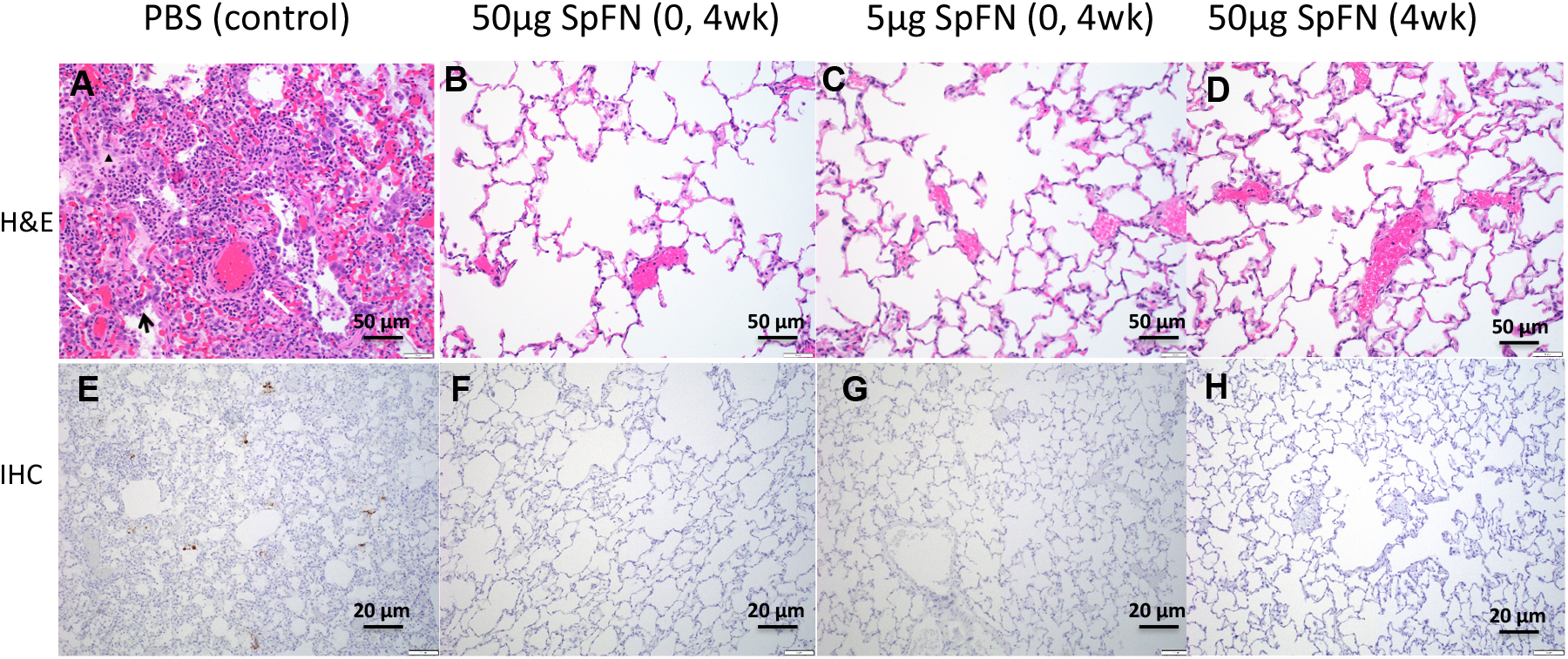
Histopathology and virus detection in the lungs of SpFN vaccinated and unvaccinated control rhesus macaques following SARS-CoV-2 respiratory challenge. At 7 days post-challenge paraffin-embedded lung parenchymal tissue sections, were (**A-D**) stained with hematoxylin and eosin (H&E) and (**E-H**) for immunohistochemistry (IHC). (**E**)Viral antigen is seen in brown aggregates. Representative images are presented at two magnifications.

Whereas sgmRNA levels reached a mean of 10^7^ copies/ml in the NP swabs of control animals at day 2 after challenge, they were undetectable in six of eight animals that received two doses of 50 µg SpFN (Fig. 3B). All animals in that group had undetectable virus from day 4 onward, while virus persisted in the control animals through day 10. Five of eight controls had high levels of sgmRNA in saliva on day 2 post-challenge, whereas virus was undetectable in all animals of the two-dose 50 µg SpFN group (Fig. 3C). Total viral load in the BAL fluid, NP swabs and saliva followed trends similar to those for sgmRNA (fig. *S7*).

#### Pathologic Efficacy

Unvaccinated control animals developed histopathologic evidence of multifocal, moderate interstitial pneumonia at 7 days after challenge (Fig. 5A). The pneumonia was characterized by type II pneumocyte hyperplasia, alveolar septal thickening, edema and necrotic debris, pulmonary macrophage infiltration and vasculitis of smaller caliber blood vessels. None of the vaccinated animals, however, had evidence of interstitial pneumonia. Immunohistochemistry revealed viral antigen in alveolar pneumocytes and pulmonary macrophages in at least one lung section of every control animal (Fig. 5E). No viral antigen was detected in any vaccinated animals (Fig. 5F-H).

**Fig. 5.**
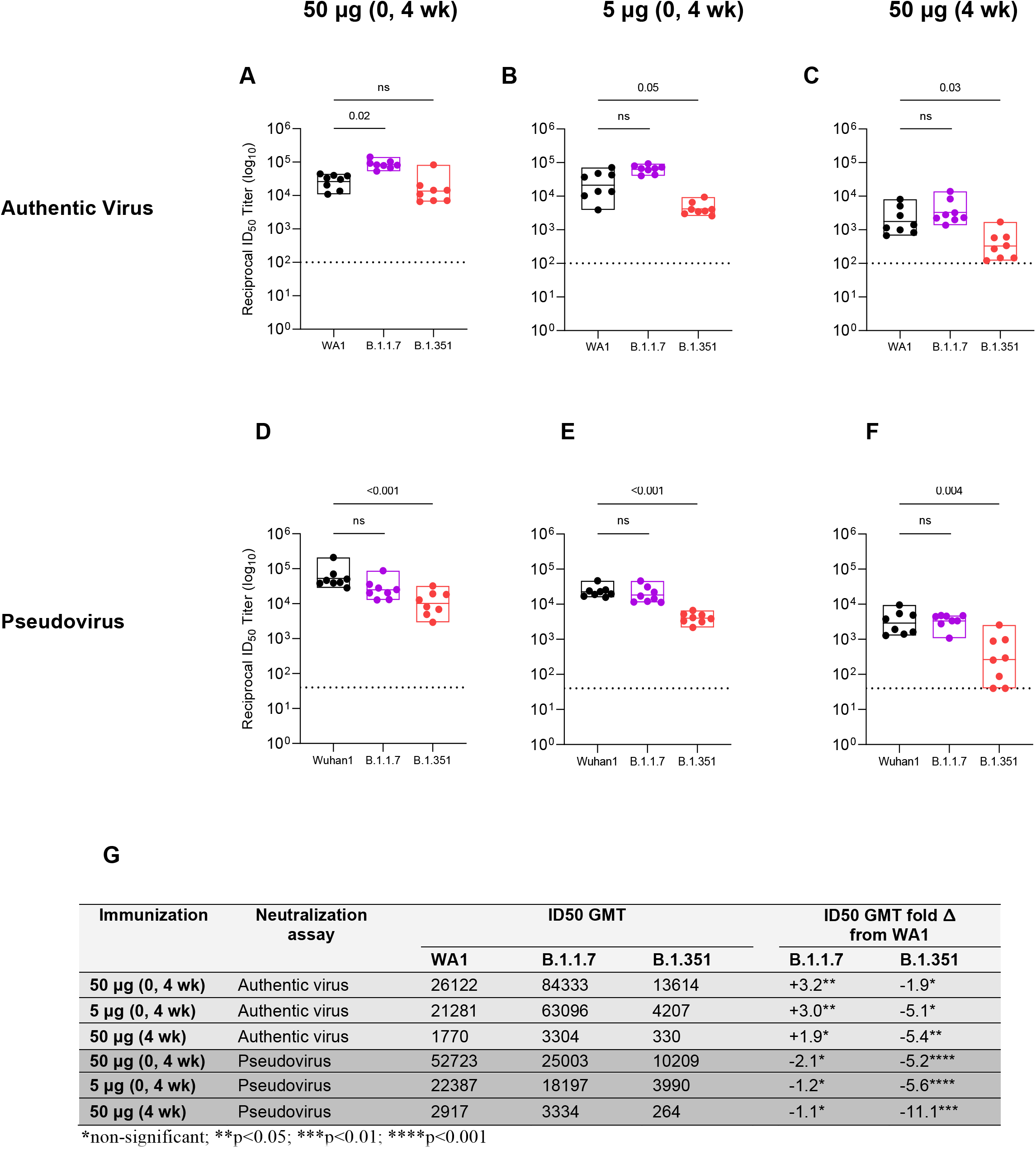
Pseudovirus and authentic virus neutralizing antibody responses elicited by SpFN vaccination in rhesus macaques against SARS-CoV-2 variants B.1.1.7 and B.1.351, as compared to responses against SARS-CoV-2 WA-1 authentic virus and Wuhan-1 pseudovirus. (**A-C**) Authentic virus and (**D-G**) pseudovirus neutralizing antibody responses were measured 2 weeks after last vaccination with either two doses of 50 µg (**A, D**) or 5 µg (**B, E**) or one dose of 50 µg (**C, F**). Reciprocal ID50 GMT fold-change from wild-type neutralization (WA-1 or Wuhan-1) was assessed (**G**), with statistical significance set at a p-value of < 0.05. Statistical comparisons were done by Kruskal-Wallis test followed by a Dunn’s post-test. In the box plots, horizontal lines indicate the mean and the top and bottom reflect the minimum and maximum.

### Breadth of Immune Response

We assessed the serum antibody responses elicited by the SpFN vaccine against two predominantly circulating SARS-CoV-2 variants of concern (VOC): B.1.1.7 and B.1.351. Serum binding assessment by biolayer interferometry to the variant forms of SARS-CoV-2 RBD showed no change in binding to B.1.1.7 (N501Y mutation) but a 25% reduction in binding to B.1.351 (K417N, E484K, N501Y mutations) in the two-dose 50 µg SpFN group, a decrement that was not statistically significant (fig. *S8*). We next assessed the serum neutralizing activity elicited by the SpFN vaccine against the two VOCs. Sera from all vaccinated NHPs elicited potent neutralizing activity against both variants in two orthogonal virus neutralization assays. Neutralization capacity of the authentic B.1.1.7 virus variant trended higher than against wild-type WA-1 across all vaccine groups (Fig. 5A-C), and was significantly so in the two-dose 50 µg group (p=0.02) (Fig. 5A). Neutralizing activity against the authentic B.1.351 virus variant, however, was diminished slightly (Fig. 5A-C), but this difference did not meet statistical significance in the two-dose 50 µg group (Fig. 5A) and was only marginally significant in the two-dose 5 µg group (p=0.05) (Fig. 5B). Neutralizing activity against the B.1.1.7 pseudovirus in the orthogonal pseudovirus assay revealed statistically equivalent ID50 GMTs to the WA-1 wild-type pseudovirus (Fig. 5D-F). Reductions in neutralizing activity against the B.1.351 pseudovirus, however, were slightly more pronounced in the pseudovirus as compared to the authentic virus assay. For example, the reciprocal ID50 GMT dropped five-fold in the two-dose 50 µg group but was still high at a value 10,209 (Fig. 5D). The absolute neutralizing antibody titers were generally elevated after two doses of vaccine, irrespective of the virus variant against which they were measured (Fig. 5G).

We expanded the assessment of the breadth of SpFN immunogenicity by interrogating the neutralizing and non-neutralizing antibody and cellular immune responses against SARS-CoV-1, Binding of vaccinee sera to SARS-CoV-1 RBD, as measured by biolayer interferometry, was absent in controls but was relatively potent in vaccinated animals—binding at half the strength of that observed to SARS-CoV-2 RBD (fig. S3, S4, Fig. 6A). Antibody-dependent cellular phagocytosis (ADCP) activity also increased remarkably in all vaccine groups, reaching a score that was 100-fold higher two-weeks after last 50 µg SpFN vaccination than baseline or unvaccinated controls (Fig. 6B). Two vaccinations with high-dose SpFN also yielded significant plaque reduction neutralizing activity against authentic SARS-CoV-1 with a reciprocal ID50 GMT of 390 (Fig. 6C) that was significantly above background (p=0.007). An orthogonal pseudovirus neutralization assay exhibited significant background activity in PBS controls. We minimized background by analyzing neutralization activity at a 90% inhibitory dilution (ID90) and found that two doses of 50 µg SpFN induced neutralizing activity against SARS-CoV-1 that was 6-fold higher (GMT 667) than controls (Fig. 6D). CD4+ T cell responses to SARS-CoV-1 S peptides, though lower in absolute percentage than to SARS-CoV-2, were still robust and strongly Th1-biased (Fig. 6E,F). The CD8+ T cell response, in contrast, was minimal (fig. S9A-C). In aggregate, the immunogenicity profile of SpFN to SARS-CoV-1, though lower in magnitude, recapitulated the quality of response observed to SARS-CoV-2.

**Fig. 6.**
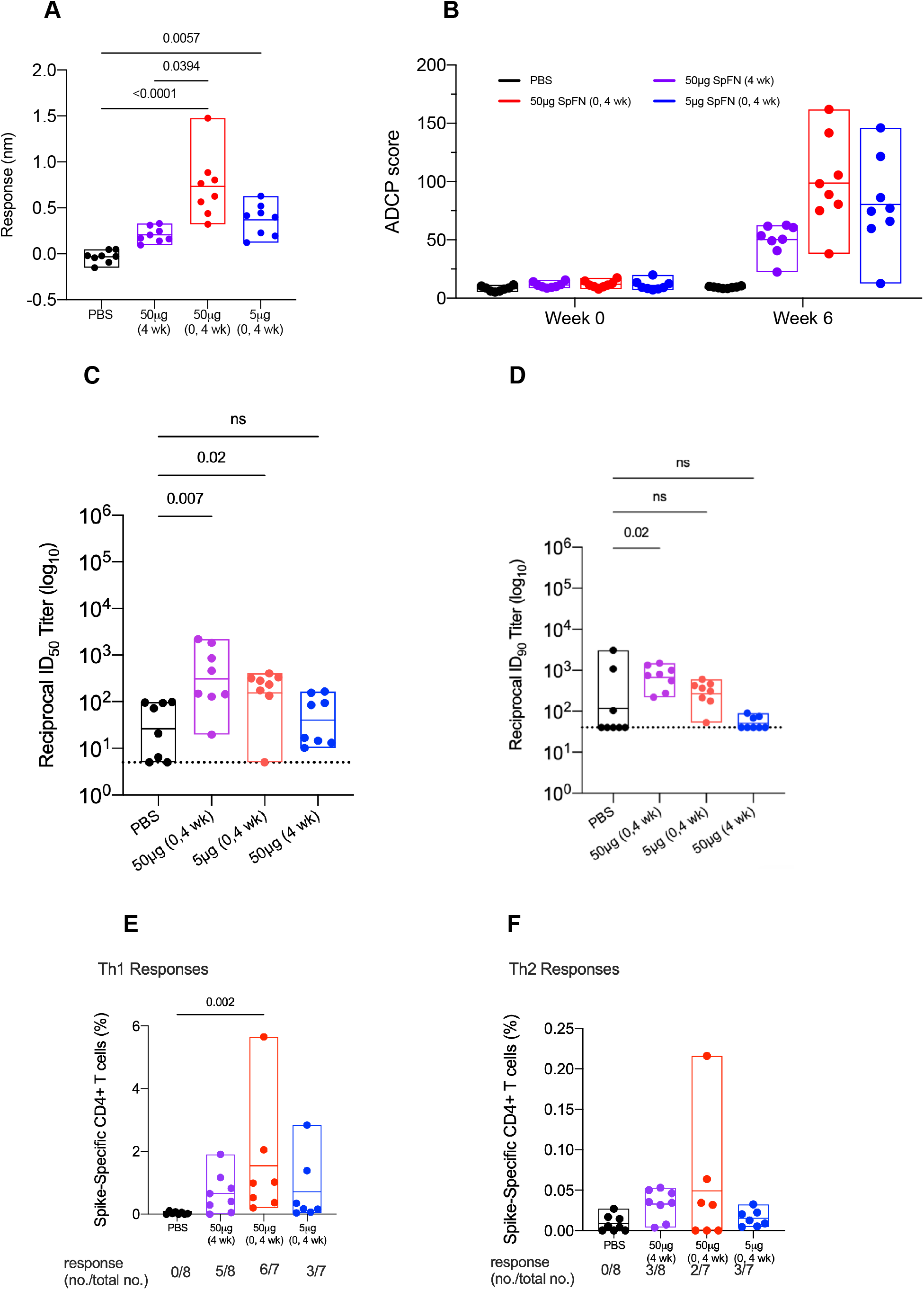
Humoral and Cellular Immune Responses to SARS-CoV-1 Elicited by SpFN in Rhesus Macaques. Serum binding responses to SARS-1 RBD by biolayer interferometry, (**B**) antibody dependent cellular phagocytosis, (**C**) authentic SARS-CoV-1 (Urbani) neutralization (ID50), (**D**) pseudo-SARS-CoV-1 (Urbani) neutralization (ID90), (**E**) SARS-CoV-1 (Urbani) Spike-specific CD4+ Th1 and (**F**) Th2 response were measured 2 weeks after last vaccination. Significance was assessed with a Kruskal-Wallis test followed by a Dunn’s post-test. In the box plots, horizontal lines indicate the mean and the top and bottom reflect the minimum and maximum.

## Discussion

The recent success in the rapid development of safe and efficacious SARS-CoV-2 vaccines has been tempered by the emergence of virus variants to which vaccine-induced immunity has shown diminished potency or efficacy (*30-34*). There remains a need, therefore, for next-generation vaccines that target the broadening antigenic diversity of SARS-CoV-2 and related coronaviruses. The major vaccines that have progressed to human efficacy trials all present SARS-CoV-2 S based on the genetic sequence of the Wuhan-Hu-1 isolate. All of these vaccines have demonstrated protective efficacy in NHPs against respiratory challenge with the closely matched USA-WA1/2020 (*25, 35-39*). These earlier animal studies, however, did not evaluate the neutralization capacity of serum against other coronavirus species. In our study of an adjuvanted SpFN vaccine, we recapitulate or surpass the protective efficacy against SARS-CoV-2 infection seen in other studies, but with a greater reduction in replicating virus against a more potent challenge than has been used previously. Additionally, we demonstrate that SpFN elicited antibodies neutralize SARS-CoV-2 ten times more potently than most vaccines and that neutralizing activity is either higher or equivalent against two major VOCs (B.1.1.7, B.1.351) in an authentic virus neutralization assay and equivalent or mildly diminished in a pseudovirus neutralization assay. Finally, SpFN demonstrates neutralization capacity of SARS-CoV-1—a separate species that has 26% and 36% sequence divergence in the S protein and S1 subunit, respectively (*40*)—above thresholds associated with protection in animals studies (*41, 42*).

SARS-CoV-2 vaccine efficacy studies in NHPs generally compare elicited antibody responses to those from recovered Covid-19 patients. We found neutralizing activity in the two-dose 50 µg SpFN group to be ten-fold higher than that in recovering patients. However, given the preponderance of animal data generated across the vaccine landscape and the lack of standardization across convalescent serum panels, we deemed it more relevant to focus immunogenicity comparisons to published data from other vaccines evaluated in NHP studies. We found that SARS-CoV-2 antibody responses in animals vaccinated with high-dose SpFN were significantly higher than those generated by high doses of leading genetic vaccines (*35, 39*). That differential increased when SpFN was compared to recombinant adenovirus vector vaccines (*25, 38*). The platform closest to SpFN in composition and design is the adjuvanted S-2P rosette, NVX-CoV2373. In a study of cynomolgus macaques, NVX-CoV2373 induced a SARS-CoV-2 antibody responses that exceeded other vaccines but was lower than that generated by SpFN. Direct quantitative comparisons of NHP immunogenicity and efficacy studies, however, can be difficult to interpret, as doses and platforms vary across studies, and immunologic and virologic endpoints are not measured by validated or identical assays. We have made attempts to overcome this limitation by analyzing specimens with orthogonal assays harmonized to consensus platforms. Additionally, our pseudovirus neutralization assay demonstrated equivalence to others in a multi-site concordance survey of reference laboratories.

Potent neutralizing antibody responses may offer advantages for both vaccine efficacy and durability. Thus far, neutralizing activity has been predictive of efficacy in human trials, as vaccines that generate lower antibody titers have diminished efficacy (*34*). An open question remains, however, regarding the length of immunity conferred by SARS-CoV-2 vaccines. For those infectious diseases that are contained by neutralizing antibodies, peak titers have been shown to predict durability (*43-45*). As such, SpFN may offer longer protection than counterparts; though this requires empirical confirmation.

Cross-neutralizing activity against SARS-CoV-2 VOCs is largely diminished for other vaccines at an approximately ten-fold reduction. SpFN induced serum cross-neutralizing responses, however, that were not significantly reduced. Additionally, we found serum binding to mutated SARS-CoV-2 RBD was either unaffected or mildly diminished. Cross-neutralizing activity against SARS-CoV-1 has not been reported yet for other vaccines in advanced development. Some early reports of nanoparticle vaccine approaches presenting RBD have begun to show breadth of neutralizing but these studies either have been limited murine immunogenicity and have yet to demonstrate large animal efficacy or do not generate a full repertoire of multi-site directed antibodies given their restriction to one domain of the S glycoprotein (*46-48*). We found a comprehensive binding and neutralizing antibody response and a balanced cellular immune response against SARS-CoV-1 Although background neutralizing activity was high in one assay, neutralizing potency against SARS-CoV-1 was confirmed in an orthogonal virus neutralization assay. Still, we are testing purified IgG from vaccine serum in both assays to control for the background levels at baseline and in controls.

We hypothesize that breadth of immune response across the SARS betacoronaviruses elicited by the SpFN vaccine may be the result of several factors. First, the quantity of the polyclonal antibody response may surpass a threshold that overcomes resistance to neutralization of antigenically distinct virus variants. Second, repetitive, ordered display of antigen on a self-assembling nanoparticle has been shown to drive an expanded germinal center reaction with resultant increases in B cell receptor mutation, affinity maturation and plasma cell differentiation (*6-8*). Lastly, the adjuvant, ALFQ, may drive some of the breadth through CD4 T cell activation (*49, 50*), especially given the high Th1 response elicited by the co-formulation. Epitope mapping and adjuvant comparison studies are underway to dissect the immune potency and breadth.

The collective immune response elicited by SpFN translated into a robust and rapid reduction in replicating virus in the upper and lower airways of animals and resultant prevention of pulmonary pathology. It is notable that SpFN protected against a viral challenge that was higher than in any other NHP study to date, as replicating virus levels detected in the upper and lower airways of unvaccinated controls reached a mean of 10^6^ to 10^7^ copies/ml. Despite this higher challenge, SpFN still protected as rapidly as other leading vaccines. SpFN also quickly eliminated sgmRNA in NP swabs, which has implications for decreasing viral shedding and transmission. The absence of an antibody titer boost after challenge suggests a potent anamnestic response and lends additional evidence for near-sterilizing immunity, which again may have implications for preventing viral transmission. Altogether, the immunologic potency and breadth and virologic and pathologic efficacy of SpFN in NHPs support its advancement to evaluation in a phase 1 clinical trial.

## ACKNOWLEDGMENTS

We thank J. Lay, E. Zografos, J. Lynch, L. Mendez-Rivera, N. Jackson, B. Silke, U. Tran and S. Peters for technical support, assistance and advice.

## Author contributions

K.M. and M.G.J designed the study. H.A.K., S.V., N.L.M., D.B. provided additional inputs into study design modifications. I.E.N, A.A., K.K.P., C.M.C., C.S., R.E.C, P.V.T., W-H.C., R.S.S., A.H., E.J.M., C.E.P., W.C.C., M.C., C.S., P.J.L., A.A., K.M.W., M.D., I.S., J.R.C., K.G.L, V.D., S.M., K.A., R.C., S.J.K. D.M.P., N.K., V.R.P., Y.H., L.L.J., G.D.G. performed immunologic and virologic assays. H.A.E, A.C., M.G.L. led the clinical care of the animals design and interpretation of data. S.P.D., X.Z., E.K.D performed histopathology. K.M., M.G.J., P.V.T., W-H.C., R.S.S., A.H., E.J.M., C.E.P., W.C.C., and M.C. designed the immunogens. M.R., G.R.M., and A.A. designed and provided the adjuvant. M.G.J., H.A.K. J.A.H., M.G.T., C.S. R.J.O., S.P.D., M.F.A., S.V., P.T.S., D.D.H., M.S.D., M.G.L., M.R., G.G.D., S.A.P., N.L.M. D.L.P. and K.M. analyzed and interpreted the data. K.M. wrote the paper with assistance from all coauthors.

## Data and materials availability

All data are available in the manuscript or the supplementary materials.

