## Supplemental Materials for "Efficacy of a Broadly Neutralizing SARS-CoV-2 Ferritin Nanoparticle Vaccine in Nonhuman Primates"

### Materials and Methods

**Vaccine and adjuvant design and production.** The Spike Ferritin Nanoparticle (SpFN) vaccine immunogen was produced by linking *Helicobacter pylori* ferritin to the C-terminal region of the pre-fusion stabilized ectodomain (residues 12-1158) of the SARS-CoV-2 S protein. The adjuvant, Army Liposomal Formulation with QS21 (ALFQ) was prepared as previously described. Briefly, ALFQ is a unilamellar liposome that contains saturated phospholipids, cholesterol, monophosphoryl lipid A and the saponin, QS-21. It is comprised of dimyristoyl phosphatidylcholine (DMPC), dimyristoyl phosphatidylglycerol (DMPG), cholesterol (Chol), and synthetic monophosphoryl lipid A (3D-PHAD®) (Avanti Polar Lipids, Alabaster, AL) and QS-21 (Desert King, San Diego, CA). ALFQ formulation has been described previously.(*1*) Briefly, DMPC and cholesterol were dissolved in chloroform and DMPG and 3D-PHAD® were dissolved in chloroform:methanol 9:1. The lipids were mixed in a molar ratio of 9:1:12.2:0.36 (DMPC:DMPG:Chol:3D-PHAD®) and the solvent was removed by rotary evaporation. 3D-PHAD® and QS-21 doses were 200 and 100 µg, respectively. The lipids were suspended in Sorenson's PBS, pH 6.2, microfluidized to form small unilamellar vesicles (SUV) and filtered. QS-21 was solubilized in Sorenson's PBS, pH 6.2, filtered and added to the SUV to form ALFQ. The final lipid ratio was 9:1:12.2:0.114:0.044 (DMPC:DMPG:Chol:3D-PHAD®:QS-21).

Immunogen was diluted in dPBS (Lot#723188, Quality Biological, Gaithersburg, MD) to 0.1 mg/ml or 0.01 mg/ml and mixed 1:1 with 2X ALFQ liposomes on a tilted slow-speed roller at room temperature for 10 min, followed by incubation at 4$^{o}$C for 50 min. All reagents were equilibrated to room temperature before use and immunizations were performed within 4 h of vaccine formulation. Each vaccine comprised a 1.0 ml solution of SpFN formulated with ALFQ.

**Study design and procedures.** Thirty-six male and female specific-pathogen-free, research-naïve Chinese-origin rhesus macaques (age 3 - 7 years) were distributed—on the basis of age, weight and sex—into 4 cohorts of 8 animals (Table S1). Animals were vaccinated intramuscularly with either 50 or 5 μg of SpFN, formulated with ALFQ, or 1ml of PBS in the anterior proximal quadriceps muscle, on alternating sides with each dose in the series. Immunizations were administered twice, 4 weeks apart, or once, 4 weeks prior to challenge. Animals were challenged with virus stock obtained through BEI Resources, NIAID, NIH: SARS-Related Coronavirus 2, Isolate USA-WA1/2020, NR-53780 (Lot# 70038893). Virus was stored at -80°C prior to use, thawed by hand and placed immediately on wet ice. Stock was diluted to $5x10^5$ $TCID_{50}$/ml in PBS and vortexed gently for 5 seconds prior to inoculation.

All procedures were carried out in accordance with institutional, local, state and national guidelines and laws governing animal research included in the Animal Welfare Act. Animal protocols and procedures were reviewed and approved by the Animal Care and Use Committee of both the US Army Medical Research and Development Command (USAMRDC, protocol 11355007.03) Animal Care and Use Review Office as well as the Institutional Animal Care and Use Committee of Bioqual, Inc. (protocol number 20-092), where nonhuman primates were housed for the duration of the study. USAMRDC and Bioqual, Inc. are both accredited by the Association for Assessment and Accreditation of

Laboratory Animal Care and are in compliance with the Animal Welfare Act and Public Health Service Policy on Humane Care and Use of Laboratory Animals.

**Serum antibody assessments.** ***Binding antibody measurements.*** SARS-CoV-2-specific binding IgG antibody responses were measured using MULTI-SPOT® 96-well plates, (Meso Scale Discovery (MSD), Rockville, MD). Multiplex wells were coated with three SARS-CoV-2 antigens, S, RBD and Nucleocapsid (N) at a concentration of 200-400 ng/ml and BSA, which served as a negative control. 4-plex MULTISPOT plates were blocked with MSD Blocker A buffer for 1 hour at room temperature (RT) while shaking at 700 rpm. Plates were washed with buffer before the addition of reference standard and calibrator controls. Serum samples were diluted at 1:1,000 - 1:100,000 in diluent buffer, then added to each of four wells. Plates were incubated for 2 hours at room temperature while shaking at 700 rpm, then washed. MSD SULFO-TAG™ anti-IgG antibody was added to each well. Plates were incubated for 1 hour at RT with shaking at 700 rpm and washed, then MSD GOLDTM Read buffer B was added to each well. Plates were read by the MESO SECTOR SQ 120 Reader. IgG concentration was calculated using DISCOVERY WORKBENCH® MSD Software and reported as arbitrary units (AU)/ml.

The ability of SARS-CoV-2 Spike-specific binding antibodies to inhibit S or RBD binding to the ACE-2 receptor was also measured using MULTI-SPOT® 96-well plates (MSD, Rockville, MD). Antigen-coated plates were blocked and washed as described above. Assay calibrator and samples were diluted at 1:25 - 1:1,000 in MSD Diluent buffer, then added to the wells. Plates were incubated for 1 hour at room temperature while shaking at 700 rpm. ACE2 protein conjugated with MSD SULFO-TAG™ was added and plates were incubated for 1 hour at room temperature while shaking at

700rpm. Plates were washed and read as described above. AU/ml concentration of inhibitory antibodies was calculated with DISCOVERY WORKBENCH® MSD Software.

Binding antibody measurements by octet biolayer interferometry were made on biosensors of the Octet FortéBio Red96 instrument (Sartorius, Fremont CA) that were hydrated in PBS prior to use. All assay steps were performed at 30°C with agitation set at 1,000 rpm. Baseline equilibration of the anti-His-tag biosensors (HIS1K biosensors with a conjugated Penta-His antibody (Sartorius, Fremont, CA) was carried out with PBS 15 s, prior to SARS-CoV2-RBD (30ug/ml diluted in PBS) loading for 120 s. After dipping in assay buffer (15 s in PBS), biosensors were dipped in the serum samples (100-fold dilution) for 180 s. Binding response (nm) at 180 s was recorded for each sample.

***Virus neutralization assessments.*** *SARS-CoV-1 and SARS-CoV-2 Pseudovirus Neutralization.* The S expression plasmid sequence for SARS-CoV-2 was codon optimized and modified to remove an 18 amino acid endoplasmic reticulum retention signal in the cytoplasmic tail to improve S incorporation into pseudovirions (PSV) and thereby enhance infectivity. SARS-CoV-2 pseudovirions (PSV) were produced by co-transfection of HEK293T/17 cells with a SARS-CoV-2 S plasmid (pcDNA3.4), derived from the Wuhan-Hu-1 genome sequence (GenBank accession number: MN908947.3) and an HIV-1 NL4-3 luciferase reporter plasmid. Infectivity and neutralization titers were determined using ACE2-expressing HEK293 target cells (Integral Molecular, Philadelphia, PA) in a semi-automated assay format using robotic liquid handling (Biomek NXp Beckman Coulter, Brea, CA). Virions pseudotyped with the vesicular stomatitis virus (VSV) G protein were used as a non-specific control. Test sera were diluted 1:40 in growth medium and serially diluted; then 25 µl/well was added to a white 96-well plate. An equal volume of diluted SARS-CoV-2 PSV was added to each well and plates were incubated for 1 hour at 37°C. Target cells were added to each well (40,000 cells/ well) and plates were incubated for an additional 48 hours. Relative light units (RLU) were measured with the EnVision Multimode Plate Reader (Perkin Elmer, Waltham, MA) using the Bright-Glo Luciferase Assay System (Promega, Madison, WI). Neutralization dose–response curves were fitted by nonlinear regression using the LabKey Server. Final titers are reported as the reciprocal of the dilution of serum necessary to achieve 50% ($ID_{50}$, 50% inhibitory dose), 80% neutralization (ID80, 80% inhibitory dose) and 90% neutralization (ID90, 90% inhibitory dose). Assay equivalency was established by participation in the SARS-CoV-2 Neutralizing Assay Concordance Survey (SNACS) run by the Virology Quality Assurance Program and External Quality Assurance Program Oversite Laboratory (EQAPOL) at the Duke Human Vaccine Institute, sponsored through programs supported by the National Institute of Allergy and Infectious Diseases, Division of AIDS.

*Authentic SARS-CoV-2 wild-type neutralization assay.* Authentic virus neutralization was measured using SARS-CoV-2 (2019-nCoV/USA_WA1/2020) that was obtained from the Centers for Disease Control and Prevention and passaged once in Vero CCL81 cells (ATCC). Rhesus sera were serially diluted and incubated with 100 focus-forming units (FFU) of SARS-CoV-2 for 1 h at 37°C. Serum-virus mixtures were added to Vero E6 cells in 96-well plates and incubated for 1 h at 37°C. Cells were overlaid with 1% (w/v) methylcellulose in MEM. After 30 h, cells were fixed with 4% PFA in PBS for 20 min at room temperature then washed and stained overnight at 4°C with 1 μg/ml of antibody CR3022 in PBS supplemented with 0.1% saponin and 0.1% bovine serum albumin. Cells were subsequently stained with HRP-conjugated goat anti-human IgG for 2 h at room temperature. SARS-CoV-2-infected cell foci were visualized with TrueBlue peroxidase substrate (KPL) and quantified using ImmunoSpot microanalyzer (Cellular Technologies). Neutralization curves were generated with Prism software (GraphPad Prism 8.0).

*Authentic SARS-CoV-2 and SARS-CoV-1 variant neutralization assay.* The SARS-CoV-2 viruses USA-WA1/2020 (WA1), USA/CA_CDC_5574/2020 (B1.1.7), and hCoV-19/South Africa/KRISP-EC-K005321/2020 (B1.351) were obtained from BEI Resources (NIAID, NIH) and propagated for one passage using Vero-E6 cells. Virus infectious titer was determined by an end-point dilution and cytopathic effect (CPE) assay on Vero-E6 cells as described previously. An end-point dilution microplate neutralization assay was performed to measure the neutralization activity of NHP serum samples. In brief, serum samples were heat inactivated and subjected to successive 3-fold dilutions starting from 1:50. Triplicates of each dilution were incubated with SARS-CoV-2 at an MOI of 0.1 in EMEM with 7.5% inactivated fetal calf serum (FCS) for 1 hour at $37^{o}$C. Post incubation, the virus-antibody mixture was transferred onto a monolayer of Vero-E6 cells grown overnight. The cells were incubated with the mixture for ~70 hours. CPE of viral infection was visually scored for each well in a blinded fashion by two independent observers. The results were then reported as percentage of neutralization at a given sample dilution.

The SARS-CoV-1 authentic virus neutralization assay methodology has been described previously (*2*). In brief, Vero E6 cells were plated on 12 well plates the day prior to use in DMEM (Dulbecco's Modified Eagle Media; Gibco, Grand Island, NY, USA) containing 10% FBS. The test serum was diluted in DMEM with 2% FBS followed by addition of the virus at ~100pfu SARS-CoV-1 (Urbani). The virus/serum suspension was incubated for 1 hour at 37C and 5%CO2. Plating medium was removed from the 12-well plates and the virus/serum suspensions were transferred to the corresponding well of the 12-well plates. The 12 well plates were then incubated for 1 hour at 37C and 5% CO2, inoculum was removed and 2mL of DMEM-2 with 20% methylcellulose was added as overlay. Plates were then incubated for 4 days at 37C and 5% CO2 before being inactivated in 10% neutral buffered formalin. Plates were stained with crystal violet for plaque enumeration.

***Antibody-Dependent Neutrophil Phagocytosis (ADNP).*** Biotinylated SARS-CoV-2 prefusion stabilized S trimer was incubated with yellow-green streptavidin-fluorescent beads (Molecular Probes, Eugene, OR) for 2 h at $37^{o}$C. 10µl of a 100-fold dilution of beads–protein was incubated for 2h at $37^{o}$C with 100µl of eight 100-fold diluted plasma samples before addition of effector cells (50,000 cells/well). Fresh human peripheral blood mononuclear cells were used as effector cells after red blood cell lysis with ACK lysing buffer (ThermoFisher Scientific, Waltham, MA). After 1h incubation at $37^{o}$C, the cells were washed, surface stained, fixed with 4% formaldehyde solution (Tousimis, Rockville, MD) and fluorescence was evaluated on an LSRII flow cytometer (BD Bioscience, San Jose, CA). Antibodies used for flow cytometry included anti-human CD3 AF700 (clone UCHT1) and anti-human CD14 APC-Cy7 (clone MϕP9) (BD Bioscience, San Jose, CA) as well as anti-human CD66b Pacific Blue (clone G10F5) (Biolegend, San Diego, CA). A phagocytic score was calculated by multiplying the percentage of bead-positive neutrophils (SSC high, CD3- CD14- CD66+) by the geometric mean of the fluorescence intensity of bead-positive cells; and dividing by 10,000.

***Antibody-Dependent Cellular Phagocytosis (ADCP).*** ADCP was measured as previously described.(*3*) Briefly, biotinylated SARS-CoV-1 or SARS-CoV-2 prefusion-stabilized S trimer was incubated with red streptavidin-fluorescent beads (Molecular Probes, Eugene, OR) for 2 h at $37^{o}$C. 10μl of a 800-fold or 900-fold (SARS-CoV-1) dilution of beads–protein was incubated for 2 h at $37^{o}$C with 100μl of eight 100-fold diluted plasma samples before addition of THP-1 cells (20,000 cells per well; Millipore Sigma, Burlington, MA). After 19 h incubation at $37^{o}$C, the cells were fixed with 2% formaldehyde solution (Tousimis, Rockville MD) and fluorescence was evaluated on an LSRII flow cytometer (BD Bioscience, San Jose, CA). The phagocytic score was calculated by multiplying the percentage of bead-positive cells by the geometric mean of the fluorescence intensity of bead-positive cells, and dividing by 10,000.

***Opsonization.*** SARS-CoV-2 Spike-expressing expi293F cells were generated by transfection with linearized plasmid (pcDNA3.1) encoding codon-optimized full-length SARS-CoV-2 Spike protein matching the amino acid sequence of the IL-CDC-IL1/2020 isolate (GenBank ACC# MN988713). Stable transfectants were single-cell sorted and selected to obtain a high-level Spike surface expressing clone (293F-Spike-S2A). SARS-CoV-2 S expressing expi293F cells were incubated with 200-fold diluted plasma samples for 30 minutes at $37^{o}$C. Cells were washed twice and stained with anti-human IgG PE, anti-human IgM Alexa Fluor 647, and anti-human IgA FITC (Southern Biotech,

Birmingham, AL). Cells were then fixed with 4% formaldehyde solution and fluorescence was evaluated on an LSRII flow cytometer (BD Bioscience, San Jose, CA).

***Antibody-Dependent Complement Deposition (ADCD).*** ADCD was performed and adapted from previously described method.(*4*) Briefly, SARS-CoV-2 S expressing expi293F cells were incubated with 10-fold diluted, heat-inactivated (56°C for 30 min) plasma samples for 30 minutes at 37°C. Cells were washed twice and resuspended in R10 media. During this time, lyophilized guinea pig complement (CL4051, Cedarlane, Burlington, Canada) was reconstituted in 1 ml cold water and centrifuged for 5 min at 4°C to remove aggregates. Cells were washed with PBS and resuspended in 200 μl of guinea pig complement, which was prepared at a 1:50 dilution in Gelatin Veronal Buffer with $Ca^{2+}$ and $Mg^{2+}$ (IBB-300x, Boston BioProducts, Ashland, MA). After incubation at 37°C for 20 min, cells were washed in PBS 15mM EDTA (ThermoFisher Scientific, Waltham, MA) and stained with an anti-Guinea Pig Complement C3 FITC (polyclonal, ThermoFisher Scientific, Waltham, MA). Cells were then fixed with 4% formaldehyde solution and fluorescence was evaluated on a LSRII flow cytometer (BD Bioscience, San Jose, CA).

***Trogocytosis.*** Trogocytosis was measured using a previously described assay.(*5*) Briefly, SARS-CoV-2 Spike expressing expi293F cells were stained with PKH26 (Sigma-Aldrich, St-Louis, MO). Cells were then washed with and resuspended in R10 media. Cells were then incubated with 200-fold diluted plasma samples for 30 min at 37°C. Effector peripheral blood mononuclear cells were next added to the R10 media at an effector to target (E:T) cell ratio of 50:1 and then incubated for 5 h at 37°C. After the incubation, cells were washed, stained with live/dead aqua fixable cell stain (Life

Technologies, Eugene, OR) and CD14 APC-Cy7 (clone MϕP9) for 15 min at room temperature, washed again, and fixed with 4% formaldehyde (Tousimis, Rockville, MD) for 15 min at room temperature. Fluorescence was evaluated on a LSRII flow cytometer (BD Bioscience, San Jose, CA). Trogocytosis was evaluated by measuring the PKH26 mean fluorescence intensity of the live CD14+ cells.

**Intracellular Cytokine Staining.** Cryopreserved peripheral blood mononuclear cells were thawed, rested for 6 h in R10 with 50U/ml Benzonase Nuclease (Sigma-Aldrich, St. Louis, MO), and stimulated with a SARS-CoV-2 S peptide pools (JPT, PM-WCPV-S) for 12 h. Cells were stained with fixable blue dead cell stain and fluorescent-labeled antibodies to cell surface markers. Sample stains were measured on a FACSymphony™ A5 SORP and analyzed on FlowJo v.9.9 software (Tree Star, Inc.). Stimulations consisted of two pools of overlapping peptides corresponding to SARS-CoV-2 S or SARS-CoV-1 S proteins (1 µg/ml, JPT, PM-WCPV-S and PM-CVHSA-S respectively) in the presence of Brefeldin A (0.65 µl/ml, GolgiPlug™, BD Cytofix/Cytoperm Kit, Cat. 555028), co-stimulatory antibodies anti-CD28 (BD Biosciences Cat. 555725 1ug/ml) and anti-CD49d (BD Biosciences Cat. 555501; 1ug/ml) and CD107a (H4A3, BD Biosciences Cat. 561348, Lot 9143920 and 253441). Following stimulation, cells were stained serially with LIVE/DEAD Fixable Blue Dead Cell Stain (ThermoFisher #L23105) and a cocktail of fluorescent-labeled antibodies (BD Biosciences unless otherwise indicated) to cell surface markers CD4-PE-Cy5.5 (S3.5, ThermoFisher #MHCD0418, Lot 2118390 and 2247858), CD8-BV570 (RPA-T8, BioLegend #301038, Lot B281322), CD45RA BUV395 (5H9, #552888, Lot 154382 and 259854), CD28 BUV737 (CD28.2, #612815, Lot 0113886), CCR7-BV650 (GO43H7, #

353234, Lot B297645 and B316676) and HLA-DR-BV480 (G46-6, # 566113, Lot 0055314). Intracellular cytokine staining was performed following fixation and permeabilization (BD Cytofix/Cytoperm, BD Biosciences) with CD3-Cy7APC (SP34-2, #557757, Lot 6140803 and 121752), CD154-Cy7PE (24-31, BioLegend # 310842, Lot B264810 and B313191), IFNγ-AF700 (B27, # 506516, Lot B187646 and B290145), TNFα-FITC (MAb11, # 554512, Lot 15360), IL-2-BV750 (MQ1-17H12, BioLegend #566361, Lot 0042313), IL-4 BB700 (MP4-25D2, Lot 0133487 and 0308726), MIP-1b (D21-1351, # 550078, Lot 9298609), CD69-ECD (TP1.55.3, Beckman Coulter # 6607110, Lot 7620070 and 7620076), IL-21-AF647 (3A3-N2.1, # 560493, Lot 9199272 and 225901), IL-13-BV421 (JES10-5A2, # 563580, Lot 9322765, 210147 and 169570) and IL-17a-BV605 (BL168, Biolegend #512326, B289357). Sample staining was measured on a FACSymphony™ A5 SORP (Becton Dickenson) and data analyzed using FlowJo v.9.9 software (Tree Star, Inc.). CD4 and CD8 T cell subsets were pre-gated on memory markers prior to assessing cytokine expression as follows: single-positive or double-negative for CD45RA and CD28. Boolean combinations of cells expressing one or more cytokines were used to calculate the sum total of Spike-specific memory CD4 or CD8 T cells. Responses from the two-peptide pools spanning SARS-CoV-2 S or SARS-CoV-1 S were summed. Display of multicomponent distributions were performed with SPICE v6.0 (NIH, Bethesda, MD).

**Total and Sub-Genomic Messenger (sgm) RNA Quantification.** Real time quantitative polymerase chain reaction was carried out for subgenomic messenger RNA (sgmRNA) and viral load RNA quantification from NP swab and BAL fluid samples. Primers targeted the envelope (E) gene of SARS-CoV-2 (Table S2). RNA was extracted from 200 µl of NP swab media or BAL specimens using the EZ1 DSP Virus kit (62724: QIAQEN) on the EZ1 Advanced XL instrument (9001874: QIAGEN). Briefly, samples were lysed in 200 µl of ATL buffer (19076: QIAGEN) and transferred to the Qiagen EZ1 for extraction. Bacteriophage MS2 (ATCC, Manassas, VA) was added to the RNA carrier and used as an extraction control to monitor efficiency of RNA extraction and amplification(*6*). Purified RNA was eluted in 90 µl elution buffer (AVE). The RT-qPCR amplification reactions were performed in separate wells of a 96-well Fast plate for the 3 targets: sgmRNA, RNA viral load, and MS2 RNA using 10ul of extracted material 0.72uM of primer and 0.2uM of probe and 1x TaqPath™ 1-Step RT-qPCR (A15299: Life Technologies, Thermo Fisher Scientific, Inc.). Amplification cycling conditions were: 2 min at 25°C, 15 min at 50°C, 2 min at 95°C and 45 cycles of 3 sec at 94°C and 30 sec at 55°C with fluorescent read at 55°C. An RNA transcript for the SARS-CoV-2 E gene was used as a calibration standard. RNA copy values were extrapolated from the standard curve and multiplied by 45 to obtain RNA copies/ml. A negative control (PBS) and two positive controls, contrived using Heat inactivated SARS-CoV-2 (ATCC, VR-1986HK), at $10^6$ and $10^3$ copies/ml, were extracted and used to assess performance of both assays.

**Histopathology.** Fresh and fixed sections of lung tissue were evaluated by light microscopy and immunohistochemistry. Lungs were perfused with 10% neutral-buffered formalin. Lung sections were processed routinely into paraffin wax, then sectioned at 5 µm, and resulting slides were stained with hematoxylin and eosin. Immunohistochemistry (IHC) was performed using the Dako Envision system (Dako Agilent Pathology Solutions, Carpinteria, CA, USA). Briefly, after deparaffinization, peroxidase blocking, and antigen retrieval, sections were covered with a mouse monoclonal anti-SARS-CoV nucleocapsid protein (#40143-MM05, Sino Biological, Chesterbrook, PA, USA) at a dilution of 1:4000 and incubated at room temperature for forty-five minutes. They were rinsed, and the peroxidase-labeled polymer (secondary antibody) was applied for thirty minutes. Slides were rinsed and a brown chromogenic substrate 3,3’ Diaminobenzidine (DAB) solution (Dako Agilent Pathology Solutions) was applied for eight minutes. The substrate-chromogen solution was rinsed off the slides, and slides were counterstained with hematoxylin and rinsed. The sections were dehydrated, cleared with Xyless, and then cover slipped. Tissue section slides were evaluated by a board-certified veterinary anatomic pathologist who was blinded to study group allocations. Immunohistochemistry (IHC) was performed with Dako Envision.

**Convalescent Plasma Samples.** A panel of 41 human convalescent-phase plasma samples were obtained from BEI Resources Repository (N=30), StemExpress (East Norritin, PA) (N=7) and a Walter Reed Army Institute of Research institutional review board-approved leukapheresis protocol (#1386H) (N=4) for which written informed consent was provided by participants. Samples were collected from males (N=20) and females (N=21) ranging in age from 31 to 71 years. Individuals donated plasma specimens approximately four-to-eight weeks after laboratory-confirmed SARS-CoV-2 infection and had histories of asymptomatic-to-mild-to-moderate clinical presentation.

**Statistical analysis.** Primary immunogenicity outputs of binding and neutralizing antibody titers as well as T cell responses were compared across vaccination groups using the Kruskal-Wallis test. Non-parametric pairwise comparisons between groups were made using the post-hoc Dunn’s test. The same hierarchical analysis was applied to comparisons of sgmRNA levels in the NP swabs and BAL fluids of vaccinated versus control groups. Statistical significance was preset at an alpha level of 0.05. The correlation between dependent T cell responses was assessed by non-parametric Spearman correlation (r).

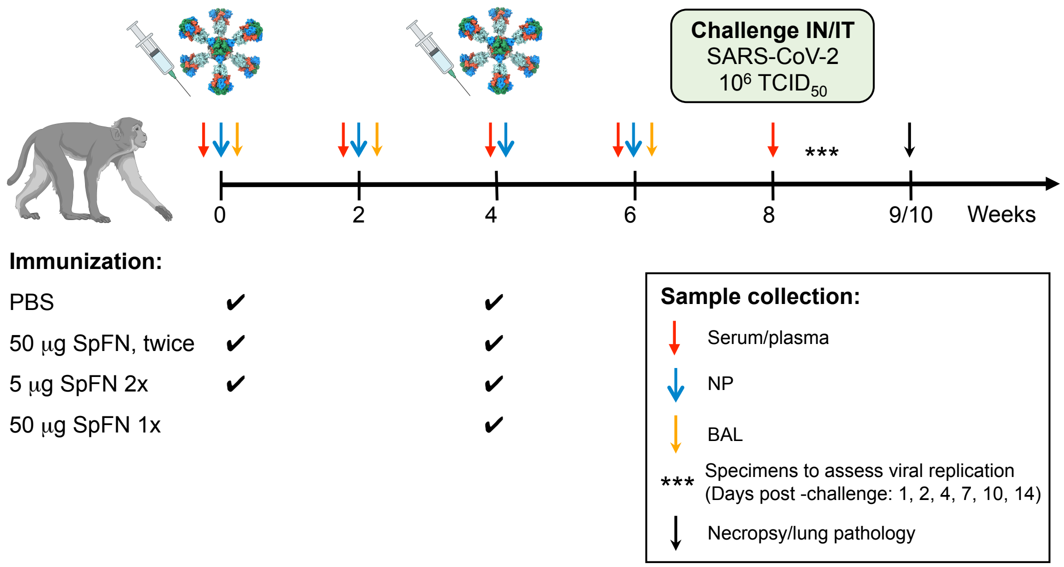


**Figure S1. Study Design and Procedures.** Thirty-six Chinese origin rhesus macaques (N=8 per group) were immunized with either 50 or 5 ug SpFN (with ALFQ) or PBS at weeks 0 and 4 or only week 4 in the one-dose 50ug group. Animals were challenged with a total of $1x10^6$ TCID50 of SARS-CoV-2 administered simultaneously by the intratracheal (1.0 ml) and intranasal (0.5 ml per nostril) route at four weeks after the last vaccination (study week 8). Serum, plasma, peripheral blood mononuclear cells, nasopharyngeal (NP) swab and bronchoalveolar lavage (BAL) fluid samples were collected every two weeks from initial vaccination through the day of challenge and stored at -80°C until analysis. Animals were followed for 7 to 8 (N=18) or 14 to 15 days (N=18) following challenge. NP swabs, saliva and BAL fluid were collected at 1, 2, 4, 7, 10 and 14 days post-challenge. Serum was collected at one-week after challenge. Pathology was performed on necropsy samples at week 9 or 10 after challenge.

**Table S1. Nonhuman primate age, sex and weight distribution by group**

| ID | Sex | Age in Years | Weight (kg) | Group | Immunogen |
|---|---|---|---|---|---|
| HS1608116 | F | 4.03 | 4.54 | 1-A | PBS |
| 171230 | F | 6.4 | 6.68 | 1-A | PBS |
| HS1605516 | F | 4.24 | 4.5 | 1-A | PBS |
| 180231 | M | 4.37 | 7.28 | 1-A | PBS |
| HS1610040 | F | 3.88 | 3.95 | 1-B | PBS |
| HS1606329 | F | 4.17 | 4.14 | 1-B | PBS |
| HS1606025 | M | 4.23 | 3.95 | 1-B | PBS |
| HS1606397 | M | 4.16 | 3.84 | 1-B | PBS |
| HS1703288 | F | 3.42 | 4.25 | 2-A | 50µg SpFN (2x) |
| HS1604100 | F | 4.39 | 4.25 | 2-A | 50µg SpFN (2x) |
| HS1606375 | M | 4.16 | 4.48 | 2-A | 50µg SpFN (2x) |
| 180262 | M | 5.45 | 8.14 | 2-A | 50µg SpFN (2x) |
| HS1607133 | F | 4.12 | 3.9 | 2-B | 50µg SpFN (2x) |
| HS1604060 | F | 4.39 | 3.95 | 2-B | 50µg SpFN (2x) |
| HS1608111 | M | 4.04 | 3.85 | 2-B | 50µg SpFN (2x) |
| HS1606395 | M | 4.16 | 3.8 | 2-B | 50µg SpFN (2x) |
| HS1705072 | F | 3.3 | 4.2 | 3-A | 5µg SpFN (2x) |
| 180203 | F | 4.38 | 5.04 | 3-A | 5µg SpFN (2x) |
| 171143 | F | 6.15 | 6.24 | 3-A | 5µg SpFN (2x) |
| 180256 | M | 4.37 | 7.2 | 3-A | 5µg SpFN (2x) |
| HS1607104 | F | 4.13 | 3.85 | 3-B | 5µg SpFN (2x) |
| HS1605136 | F | 4.3 | 3.78 | 3-B | 5µg SpFN (2x) |
| HS1606339 | M | 4.17 | 3.95 | 3-B | 5µg SpFN (2x) |
| HS1605001 | M | 4.32 | 3.85 | 3-B | 5µg SpFN (2x) |
| HS1605168 | F | 4.29 | 4.2 | 6-A | 50µg SpFN (1x) |
| 171158 | F | 5.42 | 7.78 | 6-A | 50µg SpFN (1x) |
| HS1605185 | M | 4.29 | 4.4 | 6-A | 50µg SpFN (1x) |
| 180266 | M | 4.34 | 8.56 | 6-A | 50µg SpFN (1x) |
| HS1704184 | F | 3.37 | 3.84 | 6-B | 50µg SpFN (1x) |
| HS1606076 | F | 4.22 | 3.75 | 6-B | 50µg SpFN (1x) |
| HS1608113 | M | 4.03 | 3.8 | 6-B | 50µg SpFN (1x) |
| HS1606363 | M | 4.16 | 4.15 | 6-B | 50µg SpFN (1x) |

#### Table S2. Primers and probes for SARS-CoV-2 sgmRNA and viral load

| Primer/Probe Name | Sequence 5'- 3' | Nucleotide Length |
|---|---|---|
| SARS-CoV-2 TAL E1 F | TCGTGGTATTCTTGCTAG | 18 |
| SARS-CoV-2 TAL E1 R | GAAGGTTTTACAAGACTCAC | 20 |
| SARS-CoV-2 TALE1 Probe | FAM -ACACTAGCCATCCTTACTGCG-BHQ1 | 21 |
| SARS-CoV-2 sg Leader | CGATCTCTTGTAGATCTGTTCTC | 23 |
| MS2 F | CTCTGAGAGCGGCTCTATTGG | 21 |
| MS2 R | GTTCCCTACAACGAGCCTAAATTC | 24 |
| MS2 Probe | JOE-TCAGACACGCGGTCCGCTATAACGAT- BHQ2 | 26 |

F = Forward Primer, R = Reverse Primer

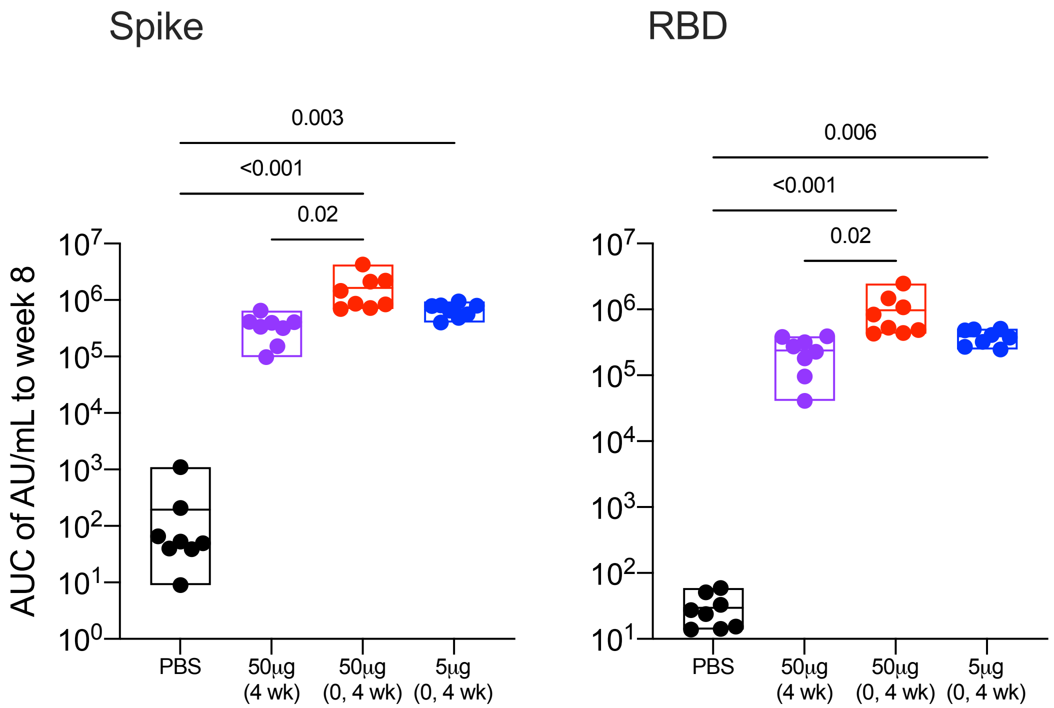


**Figure S2. IgG binding responses to prefusion stabilized spike protein and receptor binding domain by MSD electrochemiluminescence platform after SpFN vaccination in rhesus macaques** Animals were administered phosphate-buffered saline (PBS) as a control or 50µg (1 or 2 doses) or 5µg of SpFN at weeks 0 and/or 4. Serum specimens were assessed for SARS-CoV-2 Spike-specific and RBD-specific IgG by MSD (every 2 weeks following vaccination and challenge. Data shown are levels of IgG to Spike and RBD in arbitrary units (AU)/ml at 4 weeks after the last vaccination (study week 8). The horizontal line in the box plot indicates the mean; the top and bottom of the box are the minimum and maximum. Symbols represent individual animals and overlap with one another for equal values where constrained. Statistical significance of differences between groups was assessed by Kruskal-Wallis test and Dunn's post-test.

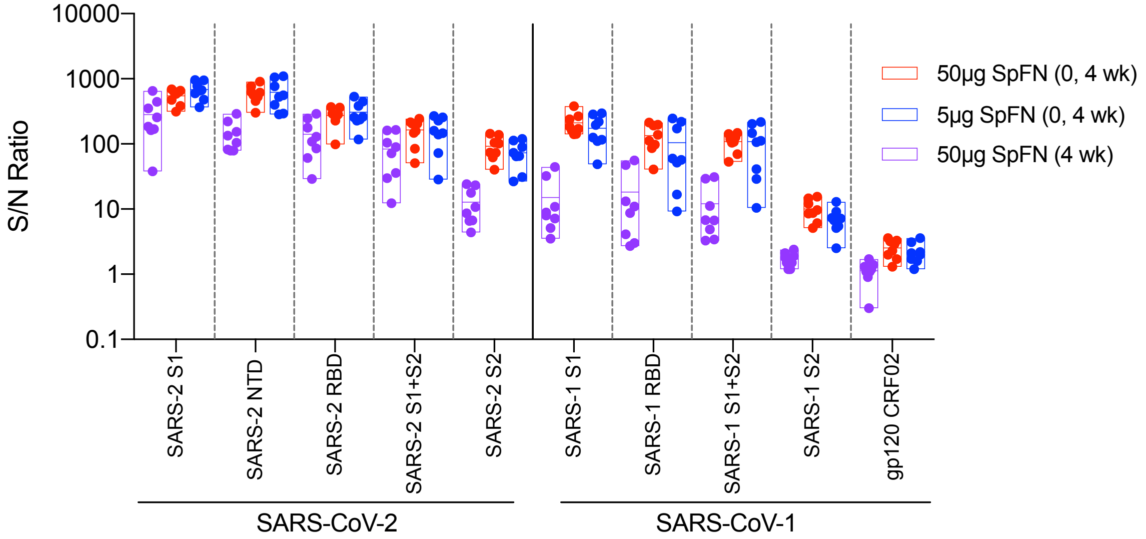


**Figure S3. Serum binding of SARS-CoV-2 and SARS-CoV antigens measured by Luminex at two weeks after last SpFN vaccination in rhesus macaques.** Plasma samples from each group were evaluated for binding to SARS-CoV-2 and SARS-CoV-1 and SARS-CoV-2 S1 and S2 subunits and receptor binding domain (RBD) and N-terminal domain (NTD) using a multiplex Luminex assay. Mean fluorescence intensity (MFI) data were divided by pre-vaccine background MFI to obtain a signal to noise (S/N) ratio of binding magnitude for each sample. Antibody binding to HIV gp120 circulating recombinant form (CRF) 02 was used as the antigen for negative control.

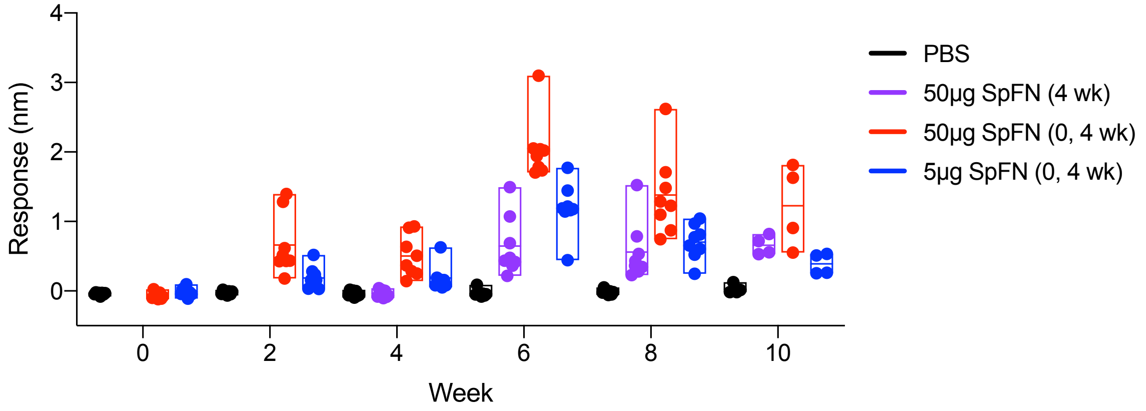


**Figure S4. Serum binding of SARS-CoV-2 receptor binding domain (RBD) measured by biolayer interferometry at two weeks after last SpFN vaccination in rhesus macaques.** SARS-CoV-2 RBD-specific binding antibody responses were assessed every 2 weeks following immunization (weeks 0, 4) and challenge (week 8).

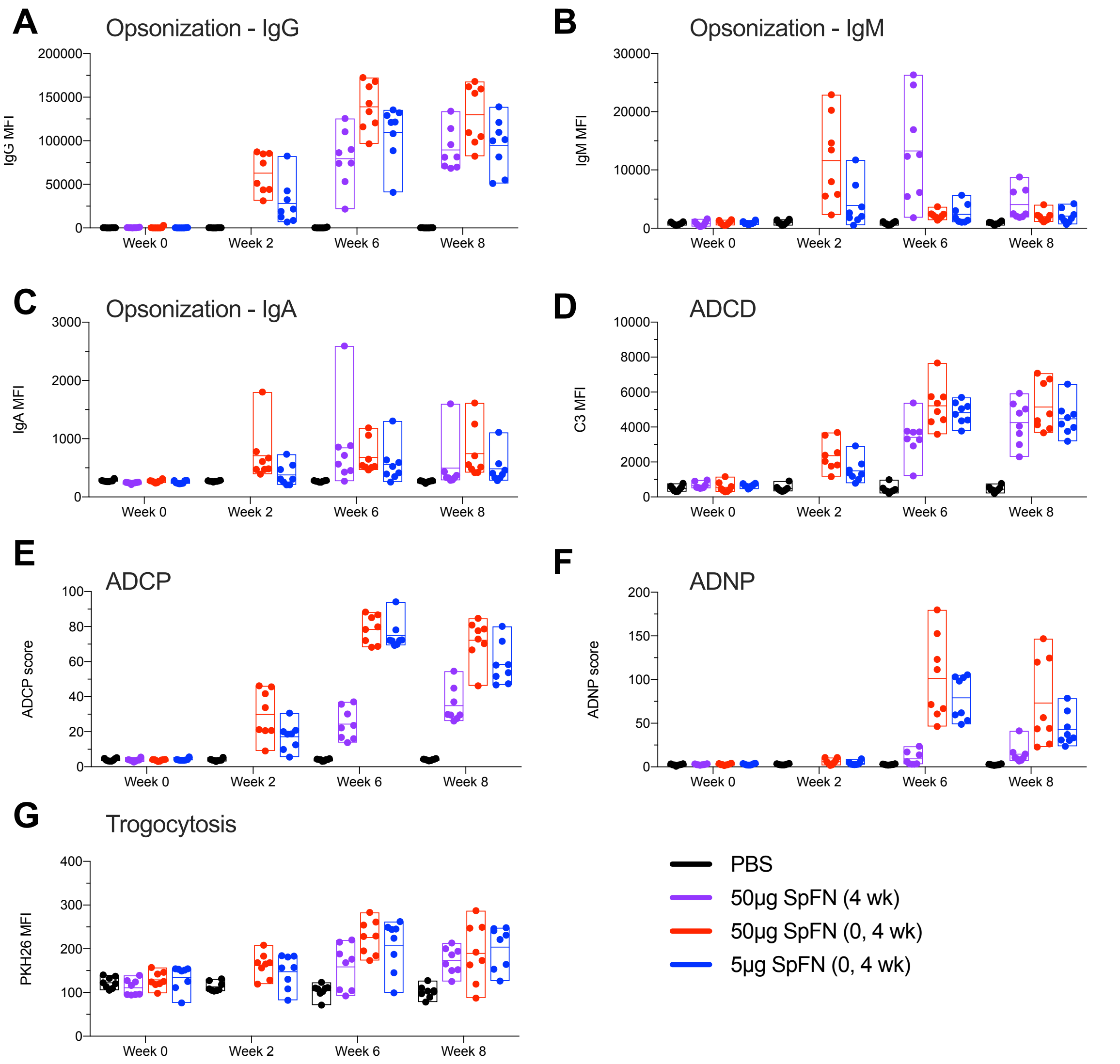


**Figure S5. Fc-mediated antibody effector responses induced by vaccination with SpFN in rhesus macaques.** Spike-expressing cells were incubated with diluted plasma from indicated time points. IgG (Panel A), IgM (Panel B), and IgA (Panel C) opsonization were measured by flow cytometry. Plasma-opsonized Spike-expressing cells were used to measure ADCD (Panel D) and Trogocytosis (Panel G). Diluted plasma was incubated with Spike-trimer coated fluorescent beads to measure ADCP (Panel E) and ADNP (Panel F).

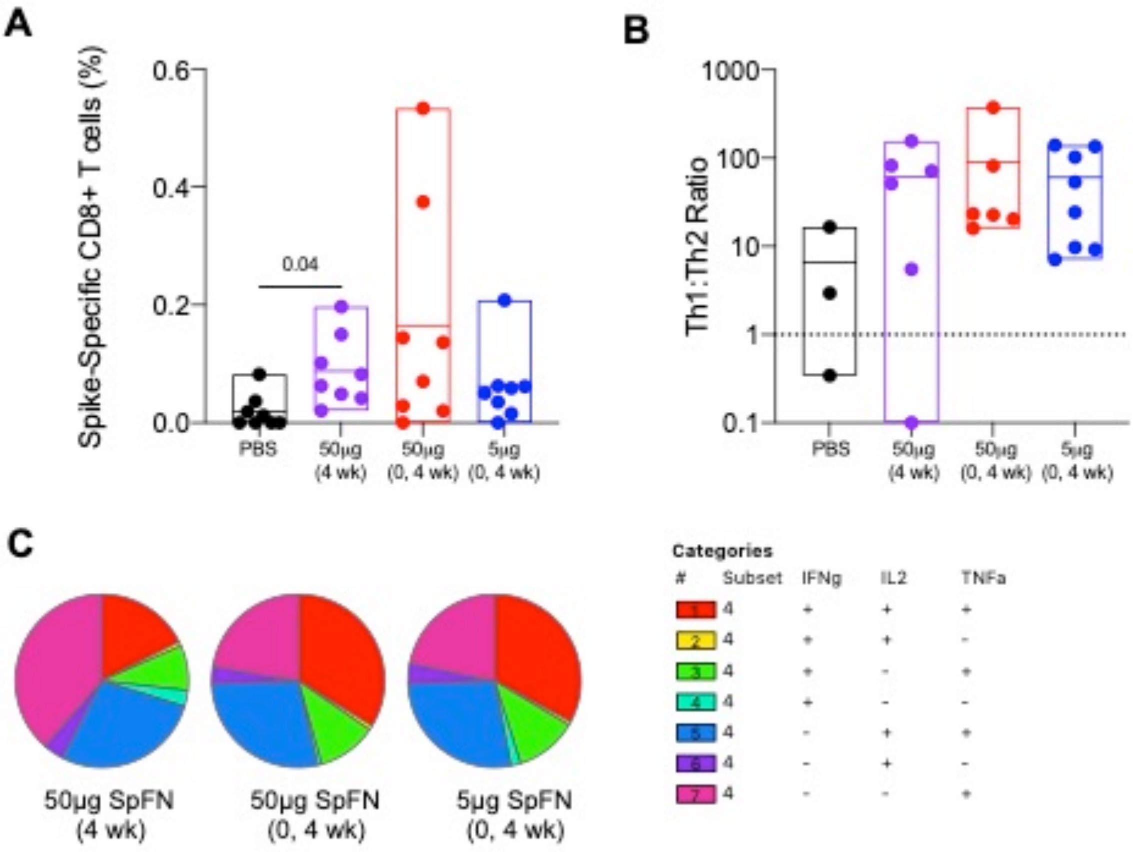


**Figure S6. CD8+ memory T cell responses and CD4+ T helper proportional responses and polyfunctionality four weeks after vaccination with SpFN in rhesus macaques.** CD8+ T cell responses shown are the summed responses from cells stimulated with Spike 1 and Spike 2 peptide pools (Panel A). Significance was assessed using a Kruskal-Wallis test followed by a Dunn's post-test. The ratio of Th1 to Th2 cells was determined at week 8 in animals with positive Th2 responses (Panel B). The dashed line indicates an equal proportion of Th1:Th2 cells. CD4+ Th1 polyfunctionality was assessed on summation of interferon gamma, tumor necrosis factor and interleukin-2 levels (Panel C).

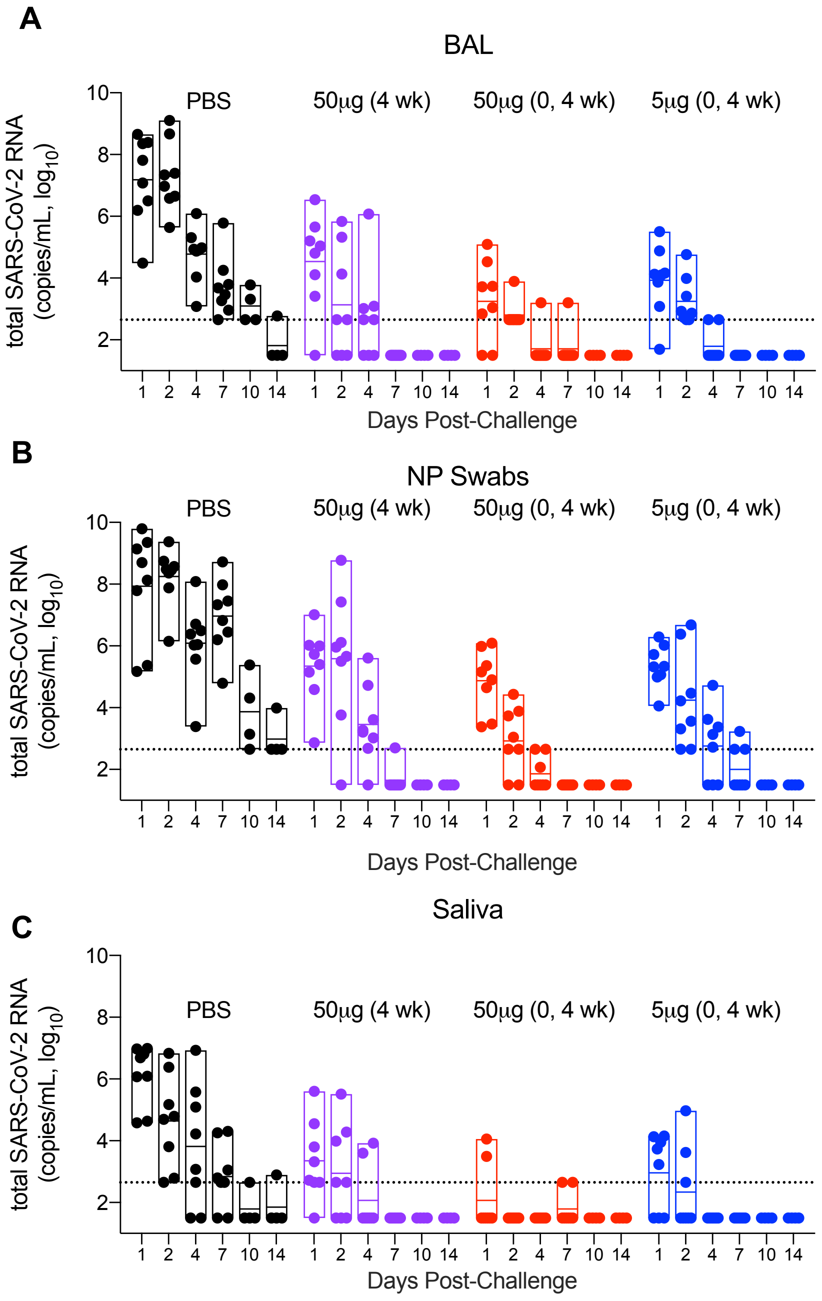


**Figure S7. Total Viral Load in the Lower and Upper Airways after SpFN Vaccination and Subsequent SARS-CoV-2 Respiratory Challenge.** Total viral RNA copies per milliliter were measured in the bronchoalveolar lavage fluid (Panel A), nasopharyngeal swabs (Panel B) and saliva of vaccinated and control animals for two weeks following intranasal and intratracheal SARS-CoV-2 (USA-WA1/2020) challenge of vaccinated and control animals. Specimens were collected on 1, 2, 4, 7, 10 and 14 days post-challenge. Dotted lines demarcate assay lower limit of linear performance range ($Log_{10}$ of 2.65 corresponding to 450 copies/ml).

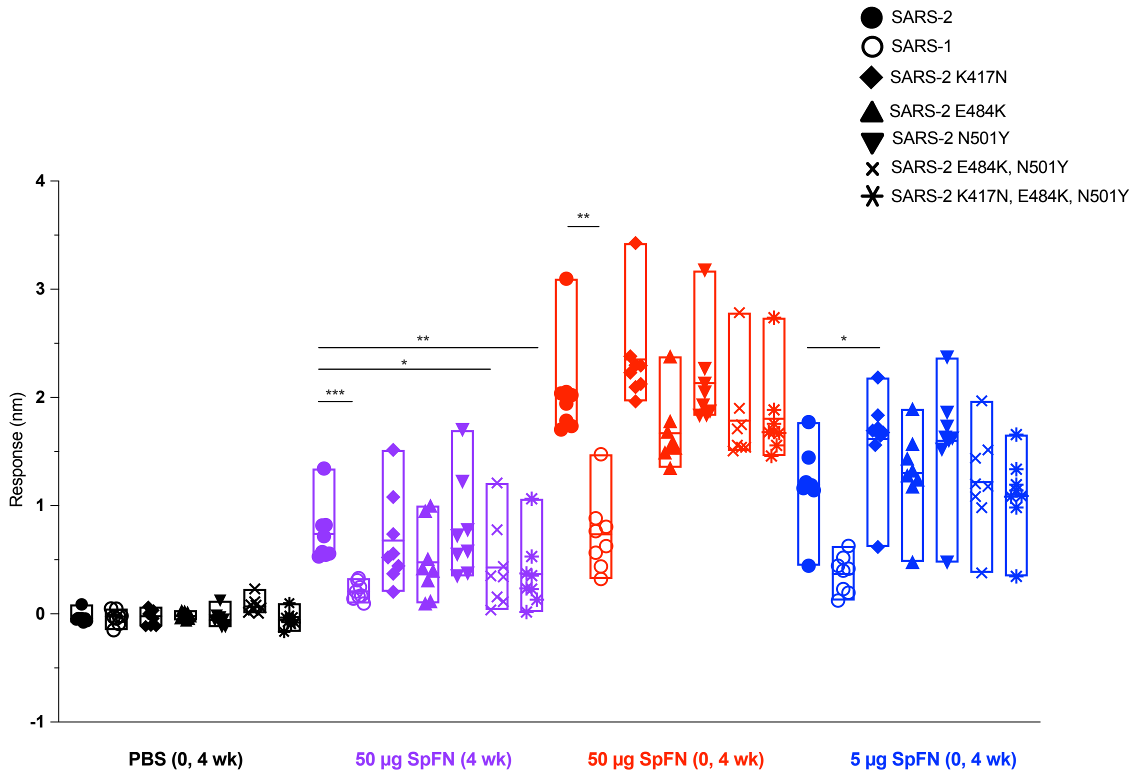


**Figure S8. Serum binding of SARS-CoV-1, SARS-CoV-2 wild type and SARS-CoV-2 mutant receptor binding domain (RBD) measured by biolayer interferometry at two weeks after last SpFN vaccination in rhesus macaques.** SARS-CoV-2 RBD-specific binding antibody responses were assessed on RBD variant forms produced by site-directed mutagenesis. Significance of differences against SARS-CoV-2 wild-type binding was assessed using a Kruskal-Wallis test followed by a Dunn's post-test. * < 0.05; ** < 0.01; *** <0.001.

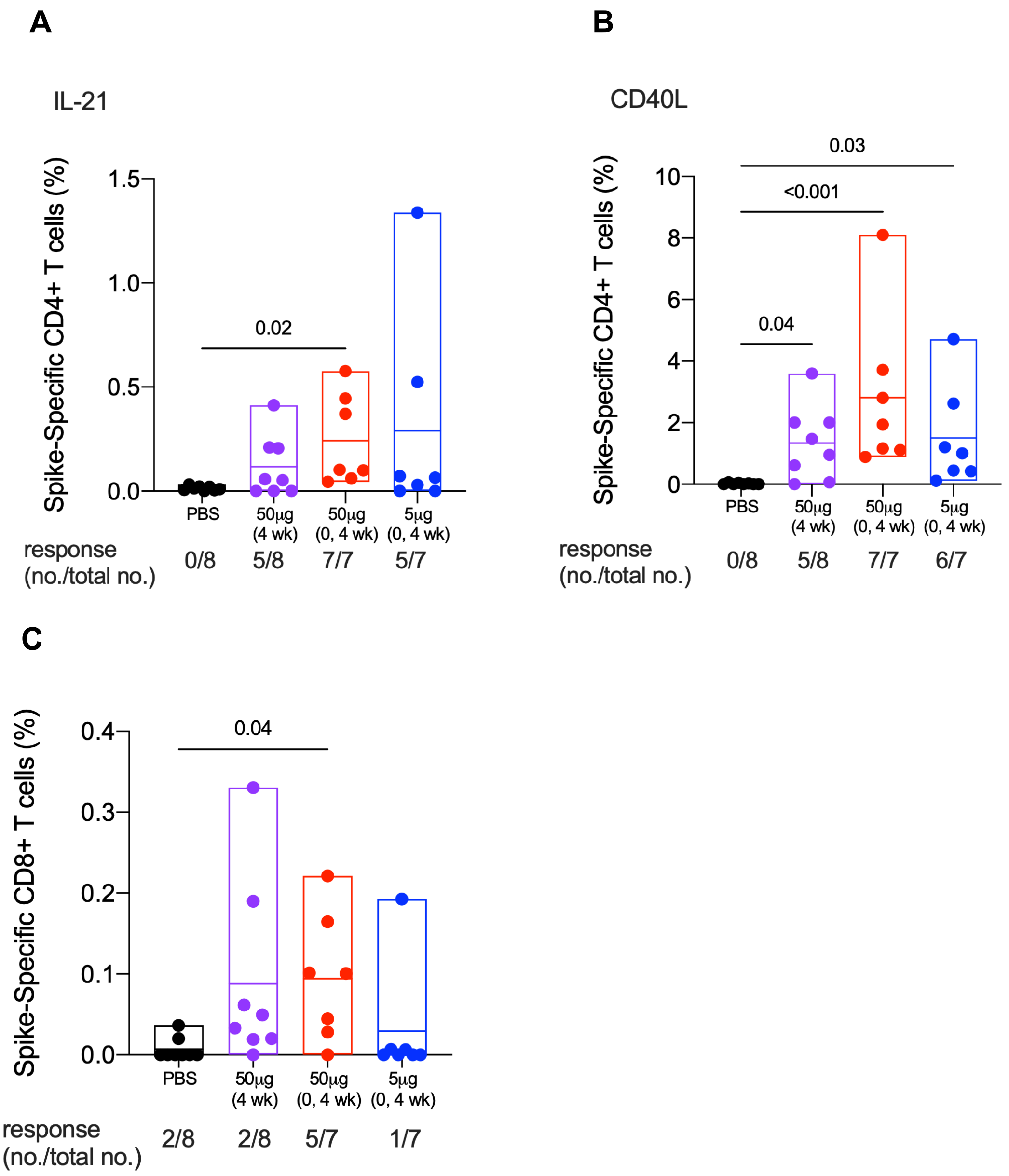


**Figure S9. Cellular immune responses elicited by SpFN vaccination in rhesus macaques against SARS-CoV-1.** T cell responses were assessed in PBMC by SARS-CoV-2 S peptide pool stimulation and ICS at 6 weeks. S-specific memory CD4+ T cells expressing the indicated marker(s) is shown as follows: (**A**) IL-21 and (**B**) CD40L. (**C**) S-specific memory CD8+ T cells expressing Th1 cytokines (IFNγ, TNF and IL-2) are shown. Boolean combinations of cytokine positive memory CD4+ T cells were summed. Probable positive responses, defined as >3 times the group background at baseline, are shown below each graph as a fraction. Significance was assessed using a Kruskal-Wallis test followed by a Dunn's post-test.
